# Obeying orders reduces vicarious brain activation towards victims’ pain

**DOI:** 10.1101/2020.06.22.164368

**Authors:** Emilie A. Caspar, Kalliopi Ioumpa, Christian Keysers, Valeria Gazzola

## Abstract

Past historical events and experimental research have shown complying with the orders from an authority has a strong impact on people’s behaviour. However, the mechanisms underlying how obeying orders influences moral behaviours remain largely unknown. Here, we test the hypothesis that when male and female humans inflict a painful stimulation to another individual, their empathic response is reduced when this action complied with the order of an experimenter (coerced condition) in comparison with being free to decide to inflict that pain (free condition). We observed that even if participants knew that the shock intensity delivered to the ‘victim’ was exactly the same during coerced and free conditions, they rated the shocks as less painful in the coerced condition. MRI results further indicated that obeying orders reduced activity associated with witnessing the shocks to the victim in the ACC, insula/IFG, TPJ, the MTG and dorsal striatum (including the caudate and the putamen) as well as neural signatures of vicarious pain in comparison with being free to decide. We also observed that participants felt less responsible and showed reduced activity in a multivariate neural guilt signature in the coerced than in the free condition, suggesting that this reduction of neural response associated with empathy could be linked to a reduction of felt responsibility and guilt. These results highlight that obeying orders has a measurable influence on how people perceive and process others’ pain. This may help explain how people’s willingness to perform moral transgressions is altered in coerced situations.

## INTRODUCTION

Many examples in the history of Mankind show that when people obey the orders from an authority, they are able to perform highly immoral acts towards others (e.g. Arendt, 1951, 1963, Herman & Chomsky, 1988). Even past experimental research, mainly by the work of Stanley Milgram (1963, 1974), showed that many people comply with coerced orders to inflict unbearable electric shock on a person for the sake of the experiment in which they were involved. However, the mechanisms underlying such drastic change in human behaviour during obedience acts remain largely unknown.

Humans, as other mammals, have the capacity to feel what others feel, namely, they have empathy. An extensive literature has shown that seeing another individual in pain triggers an empathic response in the observer (e.g. Keysers & Gazzola, 2014; Decety, 2011; Singer & Lamm, 2009; Krishnan et al., 2016). The seminal study of Singer et al. (2004) shows that experiencing painful stimulations and empathizing with the same pain delivered to others triggers an overlapping brain activity in the anterior insula (AI) and in the anterior cingulate cortex (ACC). These results, largely replicated (Fan et al., 2011; Lamm et al., 2011 for an overview), suggest that we share what others feel, at least in part, because we map their pain onto our own pain system (see Lamm & Majdandzic, 2015 for a critical review). The most widespread explanation for this phenomenon has been related to mirror neurons, which were initially shown to fire both when monkeys execute and observe an action (Gallese, Fadiga, Fogassi & Rizzolati, 1996; Keysers, 2011), but which have recently been demonstrated to also exist in the ACC (Carillo et al., 2019). It has been argued that we generally do not inflict pain to our conspecifics because we would vicariously experience this pain ourselves (Waal & Preston, 2017; Meffert et al., 2013; Smith, 1759; Hein, Morishima, Leiberg, Sul & Fehr, 2016), and we have shown that deactivating the ACC in rats, where pain mirror neurons were found, reduces the emotional reactions to the pain of others (Carrillo et al., 2019) and prevents rats from choosing actions that prevent pain to other rats over actions that harm another rat (Hernandez-Lallement et al., 2020). We therefore hypothesize that if ‘simply’ obeying the orders of an authority allows humans to perform atrocities towards other humans, it could do so by reducing the inner empathic response towards the inflicted pain, which should lead to a measurable reduction of brain response in the abovementioned regions associated with empathy and pain ratings when witnessing pain delivered under coerced compared to free condition.

Indirect evidence for this hypothesis comes from a number of studies. Caspar et al. (2016) showed that both the sense of agency and the feeling of responsibility were reduced in a condition in which people were ordered by the experimenter to inflict either a financial or a physical pain to a ‘victim’ in comparison with a condition in which they were free to decide which action to execute. Several studies further suggested that losing the sense of responsibility for an observed pain reduces activity in the neural network associated with pain empathy (Cui et al., 2015; Koban et al. 2013; Lepron et al., 2015) and reduces feelings of guilt (Yu et al., 2020). However, those studies never explored the effect of receiving orders from an authority. Cui et al. (2015) and Koban et al. (2013) focused on errors that result in pain delivered to another individual. Lepron et al. (2015) showed that when participants are the executors (vs. mere eyewitnesses in the ‘observe’ condition) of painful outcomes delivered to another individual, empathic responses (as measured by facial electromyography and heart rate variability) increased. Interestingly, the authors did not find differences in the empathic response between a condition in which they could decide and execute the action and a condition in which they could only execute an action decided by the computer. However, the perception of being “under command” may strongly differ when the instructions come from a computer and when they come from a human, authoritative figure. Therefore, whether performing an action that causes pain to others under human command would lead to reduced empathic responses to that pain is still largely unknown. Here, we predict that receiving orders to deliver painful shocks will reduce empathy for that pain, even if participants are the authors of the actions. We hypothesize that this effect could be associated with the reduced experience of responsibility and guilt under command.

In the present paradigm, one out of two volunteers (the ‘agent’) was placed in the MRI scanner and was free to decide (abbreviated as ‘Free condition’) or received a coerced instruction (‘Coerced condition’) to deliver a real, mildly painful shock to the ‘victim’ for a small monetary gain to the agent (+€0.05). We expected to observe a reduced activity of the pain network, including the insula and the ACC, as well as a reduced perception of responsibility and guilt in the coerced condition in comparison with the free condition. We also expect that participants that show more activity in these pain regions while witnessing shocks would decide to deliver fewer shocks under the free condition. These results would help explain why people are less morally inhibited when complying with orders.

Because it is difficult to associate changes in brain activity in a single location such as the anterior insula or ACC with specific mental processes such as empathy or guilt (see Lieberman & Eisenberger, 2015 vs Wager et al., 2016 for a related debate about reverse inference), we also used two multivariate signatures that have been validated to scale quite selectively with perceiving other people’s pain (the vicarious pain signature, VPS, Krishnan et al., 2016), feeling pain (the neurological pain signature, NPS, Wager et al., 2013) and with the feeling of interpersonal guilt (Yu et al., 2020).

## MATERIAL & METHOD

### Participants

Forty participants were recruited in 20 same-gender dyads. None of the participants reported to know each other. In a previous MRI study on coercion (Caspar, Beyer, Cleeremans & Haggard, under review), the sample size was 30 but involved participants who either delivered a very low number of shocks or a very high number of shocks. Given that the main contrast of interest here involved the comparison between shock and no shocks trials, we increased the sample size to 40 in order to ensure we would have enough participants delivering enough shocks to reliably estimate brain activity in both shock and no shock conditions. A priori exclusion criteria were based on the fMRI screening and head motions below 3mm. Behavioral data were not considered a priori exclusion criteria but only a post exclusion criterion. The full dataset of three participants was excluded from all the analyses: Two participants systematically disobeyed orders (one never administered shocks even when requested to do so, and one administered shocks even when requested not to do so) and one participant was not taken into account because of a late arrival. For the remaining 37 participants (11 were males), the mean age was 24.8 years old (SD=3.99). Using a voxel-wise *p*<0.001 threshold, this sample size provides 80% power to detect effects of d=0.69 in our one-tailed matched pair t-test of interest at the second level. The study was approved by the local ethics committee of the University of Amsterdam (project number: 2017-EXT-8298). Data are made available on OSF (DOI 10.17605/OSF.IO/JSHPE - direct link: https://osf.io/jshpe/).

### Procedure and material

Upon arrival in the laboratory, both participants received instructions about the experiment and provided informed consent together, ensuring that they were each aware of the other’s consent. Then, their individual pain threshold for the electrical stimulation was determined, as described in Caspar et al. (2016). Two electrodes were placed on the participants’ left hand on the abductor pollicis muscle in order to produce a clear and visible muscle twitch and the threshold was increased by steps of 1mA until a mildly painful stimulation was achieved.

By picking randomly a card in a box, participants were assigned to start either as agent or victim, but were offered the possibility to change if they wanted to. Two participants asked to start as ‘victims’ first. Participants were then told that roles would be reversed mid-way through the experiment, making the procedure fully reciprocal (full instructions are provided in ***Supplementary Material S1***). The participant who was in the role of the agent was brought into the scanner to perform the task, while the participant who was in the role of the ‘victim’ was seated nearby the control room at a table (see **Fig. 1A**). Participants in the role of the ‘victim’ were asked to place their hand on a black sheet positioned in the field of view of the camera and asked not to move her hand during the entire scanning session. The victim was invited to watch a neutral documentary to make the time pass. That neutral documentary was Planet Earth. Participants could select the documentary about Islands or the one about Mountains.

**Fig. 1.**
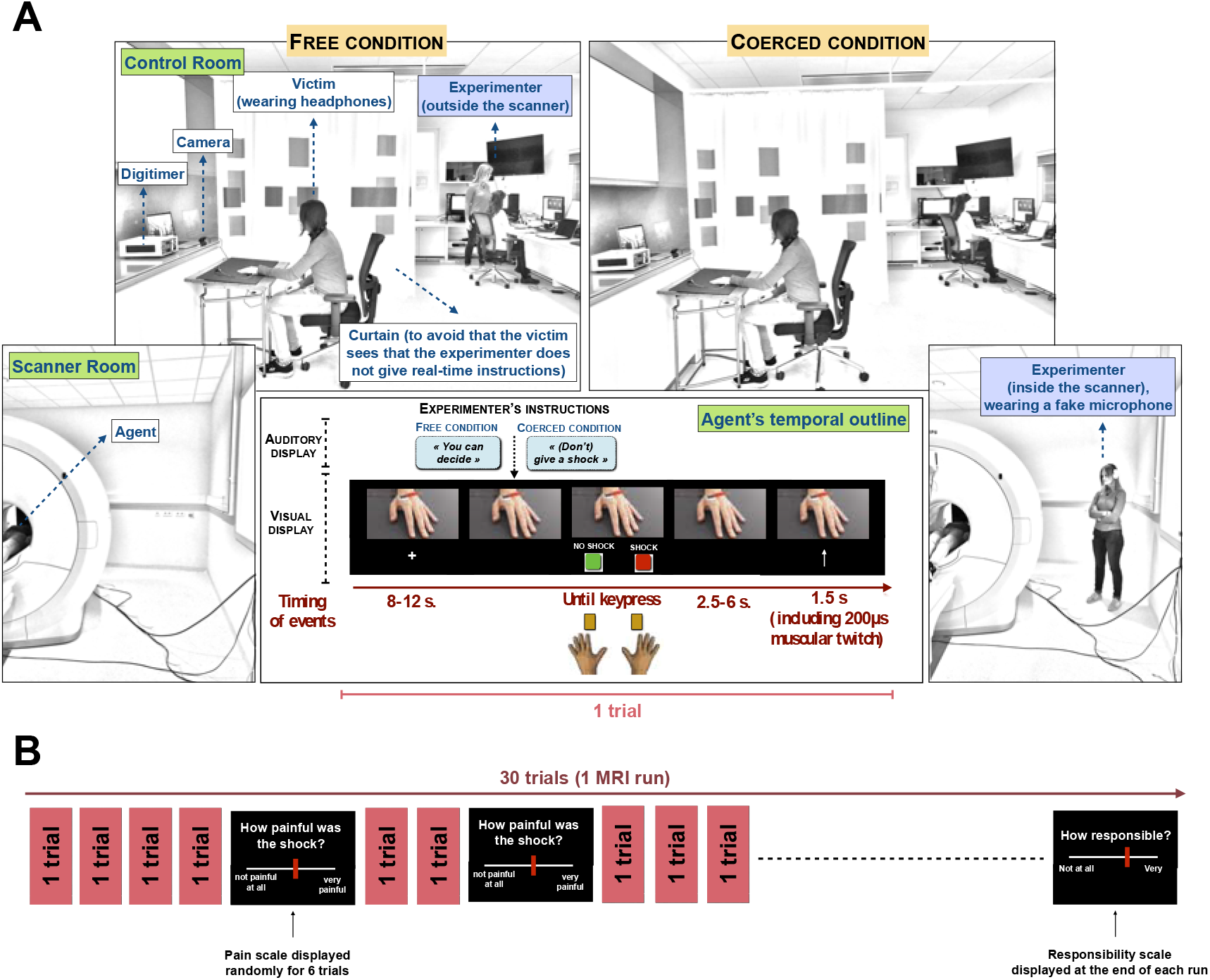
**A)** Display of the experimental set-up. The agent was inside the scanner while the ‘victim’ was outside the scanner, watching a documentary. In free blocks, the experimenter was in the control room. In coerced blocks, the experimenter entered the scanner room, indicated her presence to the agent but remained hidden from her view. **B)** Display of one single MRI run.

Each trial started with a fixation cross lasting 8-12 seconds. Then, agents heard a verbal instruction from the experimenter, in both the free and the coerced conditions. In the free condition, the experimenter told the agent ‘*you can decide*’, while in the coerced condition the experimenter said either ‘*give a shock*’ or ‘*don’t give a shock*’. In reality, these sentences were pre-recorded in order to keep control on the precise timing of each event during the scanning session. However, to increase the authenticity of the procedure, each sentence was recorded 4 times with small variations in the voice and displayed randomly. In addition, the audio recordings included a background sound similar to interphone communications. The sound of the pre-recorded instructions emitted by the computer in the control room was turned on during the free condition and turned off during the coerced condition. In case the victims (accidentally) dropped their headphones while watching the documentary, they could still believe that the experimenter gave real-time oral instructions from the control room in the free condition and from inside the scanner in the coerced condition. The sequence of orders to shock or not the ‘victim’ given by the experimenter was fully random and differed between participants.

After receiving the verbal order, a picture of two rectangles, a red one labelled ‘SHOCK’ and a green one labelled ‘NO SHOCK’, was displayed on the left and right bottom of the screen. The key-outcome mapping varied randomly on a trial-wise basis but the outcome was always fully congruent with the mapping seen by the participant. Agents could then press one of the two buttons. Pressing the SHOCK key delivered a shock to the victim while pressing the NO SHOCK key did not deliver any shocks. The shock was delivered between 2.5 and 6 seconds after the keypress (Cui et al., 2015) in order to avoid a transfer of signal between brain activity associated with the button press and brain activity associated with the visualisation of pain. Throughout the experiment, agents could observe the receiver’s hand through a real-time video displayed on the top of the screen, with electric shocks eliciting a visible muscle twitch. Seven hundred and fifty ms before the display of the shock, an arrow pointing to the top was displayed to remind participants to look at the video, even if the NO SHOCK key had been pressed. That arrow disappeared 750 ms after. This allowed the comparison of shock and no shock trials. To ensure that agents were actually watching the hand, on 6 trials in each MRI run a pain rating scale appeared, ranging from ‘*not painful at all*’ (‘0’) to ‘*very painful*’ (‘1,000’). Participants were asked to rate the intensity of the shocks seen by moving the red bar along the scale using four buttons. The keys below the middle fingers allowed participants to modify the number associated with the position of the rectangle by steps of +− 100. The keys below the index fingers allowed participants to modify the answer by steps of +− 1. After a fixed duration of 6 seconds, their answer was saved and the next trial started. If no shocks were delivered on that trial, they were asked to report that the shock was ‘*not painful at all*’. They had 6 seconds to provide their answer.

The task was split up into 4 blocks of 30 trials each, 2 blocks free choice and 2 blocks coercion (presented in 4 separate *f*MRI acquisition runs). Anatomical images were recorded between the second and third run of *f*MRI acquisition. At the end of each task block, participants rated their explicit sense of responsibility over the outcomes of their actions on an analogue scale presented on the screen, ranging from ‘*not responsible at all*’ to ‘*fully responsible*’. Each delivered shock was rewarded with €0.05 to ensure that reward prediction was similar in the two experimental conditions. In coerced blocks, participants were instructed to deliver a shock on 50% of trials, see **Fig. 1B**.

To increase the psychological effect of receiving orders, the experimenter was present in the scanner room during the coerced condition. In order to justify that the agent was still able to hear clearly the experimenter giving orders even when she was in the noisy scanner room, a fake microphone was used. The experimenter made it clear that she was present at the beginning of the coerced condition by speaking with the agent, but then moved in the corner of the room to avoid visual interference due to her presence.

At the end of the experimental session, participants were asked to fill in the Interpersonal Reactivity Index (IRI, Davis, 1980) and another questionnaire assessing what they felt during the experiment (see ***Supplementary Material S2***). Participants were paid separately, based on their own gain during the experiment.

### General data analyses

Each result was analysed with both frequentist statistics and Bayesian statistics (Dienes, 2011), except for voxel-wise brain analyses that were only analysed using frequentist approaches. Bayesian statistics assess the likelihood of the data under both the null and the alternative hypothesis. In most cases, we report BF_10_, which corresponds to the *p*(data|*H*_1_)/*p*(data|*H*_0_). Generally, a BF between 1/3 and 3 indicates that the data is similarly likely under the H1 and H0, and that the data thus does not adjudicate which is more likely. A BF_10_ below 1/3 or above 3 is interpreted as supporting H_0_ and H_1_, respectively. For instance, BF_10_=20 would mean that the data are 20 times more likely under H_1_ than H_0_ providing very strong support for H_1_, while BF_10_=.05 would mean that the data are 20 times more likely under H_0_ than H_1_ providing very strong support for H_0_ (Marsman & Wagenmakers, 2017). BF and p values were calculated using JASP (JASP Team, 2019) and the default priors implemented in JASP (Keysers, Gazzola, & Wagenmakers, 2020). Default priors used in JASP depend on the statistical tests performed (for ANOVA, see Rouder, Morey, Speckman, & Province, 2012; for t-tests, see Jeffreys, 1961; for correlations, see Jeffreys, 1961 and Ly, Verhagen, & Wagenmakers, 2016). In cases where a one-tailed hypothesis was tested, the directionality of the hypothesized effect is indicated as a subscript to the BF (e.g. BF_+0_ for a positive effect, BF_−0_ for a negative effect).

### Functional Magnetic Resonance Imagery (fMRI)

MRI images were recorded using a 3-Tesla Philips Ingenia CX system and a 32-channel head coil. T1-weighted structural images were recorded with the following specifications: matrix = 240×222; 170 slices; voxel size = 1×1×1mm. Four runs of functional images were recorded (matrix M × P: 80 × 78; 32 transversal slices in ascending order; TR = 1.7 seconds; TE = 27.6ms; flip angle: 72.90°; voxel size = 3×3×3mm, including a .349mm slice gap). Images were acquired in ascending order.

### General fMRI Data processing and first level contrasts

MRI data processing was carried out in SPM12 (Ashburner et al., 2014). EPI images were slice-time corrected to the middle slice and realigned to the mean EPI image. High quality *T*_1_ images were coregistered to the mean EPI image and segmented. The normalization parameters computed during the segmentation were used to normalize the gray matter segment (1mm×1mm×1mm) and the EPIs (2mm×2mm×2mm) to the MNI templates. Afterwards, images were smoothed with a 6mm kernel.

At the first level, we defined separate regressors for Shock and NoShock trials, with the choice and coerced trials modelled in separate runs in order to identify the activations associated with witnessing pain. Each of these regressors started 750 ms before the moment of the shock, which lasted 250 ms, up to 500 ms after the moment of the shock. This moment corresponded to when the arrow pointing to the video feedback appeared, to remind participants to watch the screen displaying the victim’s hand. The same 1.500ms-time window was taken for Shock and NoShock trials. Additional regressors included: (1) The auditory orders from the experimenter (starting between 8-12s after the start of the trial) together with the button presses (participants could press the key whenever they wanted right after the auditory orders), (2) the pain rating scale (appearing on 6/30 trials randomly 1s after the arrow pointing towards the video feedback disappeared) together with the responsibility rating scale (appearing at the end of each MRI run, 1s after the arrow pointing towards the video feedback disappeared or again 1s after the pain scale). Trials where participants disobeyed were modelled in additional regressors of no interest separately for ‘prosocial’ disobedience (i.e., they refused to administer a shock while having been ordered to send a shock) and ‘antisocial’ disobedience (i.e., they administered a shock while having been ordered not to send a shock). Finally, 6 additional regressors of no interest were included to model head translations and rotations.

At the first level, we defined the main contrast of interest to understand how receiving orders affects pain processing of the victim’s pain in comparison with not receiving orders: [FreeShock-FreeNoShock - (CoercedShock-CoercedNoShock)]. We included the contrast of Shock-NoShock rather than examining the shock condition alone in each condition to isolate the effect of witnessing a shock from carry-over activity associated with pressing the response button and seeing the arrow presented during the feedback period. To be noted, some participants administered few shocks. Because the reliability with which brain activity in the Shock condition can be estimated in the *f*MRI analysis depends on the number of trials included, only including participants delivering a large number of shocks would be ideal. However, for our results to be representative of the population, excluding too many participants delivering small numbers of shocks would bias our results towards less considerate participants. Given the distribution of shocks (i.e., {1,2,2,3,5,8,9,9,…}, see **Fig. 2**), we thus chose to only exclude participants that had delivered 1, 2, 3 or 5 shocks, and retained all participants that had delivered 8 shocks or more. This decision was not based on a power-analysis but by a subject cost-benefit analysis that pitched the benefits of inclusiveness against the cost of less stable parameter estimates. The five excluded participants were thus not included in any neuroimaging analyses.

**Fig. 2.**
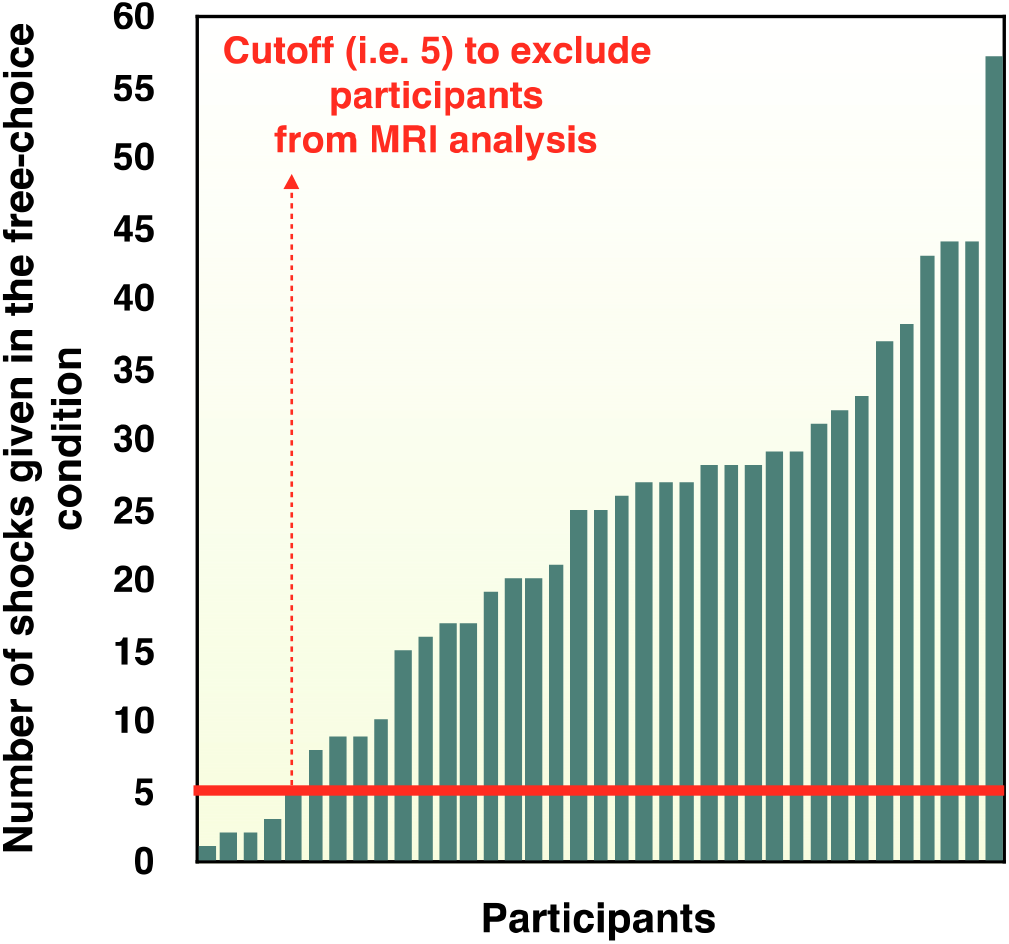
Graph with the number of shocks delivered to the victim in the free condition for each participant in increasing order. The red line represents the cut-off to include (above the red line) or exclude (below or at the level of the red line) those participants from our MRI analyses.

In order to test for the collinearity of the regressors, that is, ensuring that the brain activations observed when participants witnessed a shock or no shock on the victim’s hand are not the consequence of previous brain activations coming from anterior events, we calculated from the design matrix a Pearson correlation between our contrast of interest the [FreeShock+CoercedShock - (FreeNoShock+CoercedNoShock)] regressor and the delivery of the orders regressor. We performed correlations for each participant and obtained a mean r value of −0.056, indicating that only 0.31% (−0.056^2^*100) of the variance in the predictors is shared. We then transformed the r value of each participant into a z score and conducted a one sample t-test with 0 as the test value. The results were in favour of H_0_ (t_(36)_=−0.529, *p* = .6, Cohen’s d=−0.087 BF_10_=0.202), thus supporting the lack of influence of the order/button press regressor on the [FreeShock+CoercedShock - (FreeNoShock+CoercedNoShock)] contrast. The same analysis on the rating scale as a regressor also showed no influence on the [FreeShock+CoercedShock - FreeNoShock+CoercedNoShock] contrast (mean r = −0.022, t_(36)_=−1.159, *p* > .2, Cohen’s d = −.191, BF_10_=0.328). Thus, those results confirmed that brain activations observed during the Shock - NoShock regressor are independent from brain activations associated with the other regressors.

At the second level, we localized where this contrast [FreeShock-FreeNoShock - (CoercedShock-CoercedNoShock)] was significant across participants in a random effect one-sample t-test. Results were thresholded at *p*_*unc*_ < .001 and 5% family-wised error (FWE) corrected at the cluster level.

To decode vicarious pain intensity, we additionally used the vicarious pain signature of Krishnan et al. (2016). We converted the pattern into our image space using the ImCalc function of SPM, and then dot-multiplied each participant’s contrast [FreeShock-FreeNoShock] and [CoercedShock-CoercedNoShock] separately by this pattern. This generated a single scalar vicarious pain value for each of the 32 participants that had enough shock trials in the free condition to perform the analysis, and for each condition. The vicarious pain values were then compared using a one-tailed t-test to test our one-tailed hypothesis that vicarious pain should be higher in the free compared to the coerced condition. We used the pattern without occipital lobe, as used in Krishnan et al., to reduce visual confounds. We performed the same analysis with the neurological pain signature, NPS, Wager et al., 2013) and with the guilt-related brain signature (Yu et al., 2020).

To shed further light on the properties of the clusters resulting from the contrast [FreeShock-FreeNoShock - (CoercedShock-CoercedNoShock)] we used Marsbar (Brett, Anton, Valabregue, & Poline, 2002) to build regions of interest (ROIs) from the resulting five clusters (Table 1). With Marsbar, we computed the average signal of all the voxels in each ROI to estimate the same GLM described in our first-level, voxel-wise analysis. For each of the five ROIs, we estimated parameters of interest for the following two contrasts: [FreeShock - FreeNoShock] and [FreeShock+CoercedShock - (FreeNoShock+CoercedNoShock)]. To test our hypothesis that activity in these ROIs during shock observation is related to the decision to deliver shocks or not, we extracted the contrast values from the [FreeShock - FreeNoShock] contrast for each ROI and correlated (Pearson) them with the number of shocks freely delivered by agents. We used the [FreeShock - FreeNoShock] contrast because this is the only experimental condition in which participants could freely decide to deliver a shock or not to the victim, thus allowing us to assess the association between brain activity and decision-making. Those correlations were performed separately amongst participants that were agents first and those who were agents second. The reason for separating these two groups is because former studies indicate that if agents have been victims first, the number of shocks they have received as victims influences their decision-making (e.g., Caspar et al., 2016; 2017). Hence, one group of participants (those that had been victims first) used criteria for their decisions that differ from those of the other participants and we have reasons to believe that the decision-making process might be different enough to merit separate examination. The [FreeShock+CoercedShock - (FreeNoShock+CoercedNoShock)] contrast was established in order to obtain a general ‘pain response’, irrespective from the experimental condition. We checked with exploratory multiple linear regressions if this general effect of observing the pain experienced by another individual was associated with trait measures of empathy from the IRI subscales most associated with empathy (subscales EC, PT and PD). We excluded the FS subscale because it captures participants’ tendency to identify with fictional characters in novels and movies, and was less directly relevant (Davis 1981). This allowed us to evaluate the overall reaction of participants when they witnessed pain, no matter the experimental condition.

**Table 1.** [FreeShock-FreeNoShock - (CoercedShock-CoercedNoShock)]

| *cluster size and label*<br>*(N voxels)* | *#Voxels in cyto* | *% Cluster* | *Hem* | *Cyto or Anatomical description* | *% Area* | *cluster peak t-value* | *MNI coordinates (X,Y,Z)* |
| --- | --- | --- | --- | --- | --- | --- | --- |
| 388<br>(ROI 1: ACC) | 23.9 | 6.2 | L | Area 33 | 10.4 |  |  |
|  | 16.8 | 4.3 | R | Area 33 | 7.4 |  |  |
|  | 32.8 | 8.5 | L | Anterior Cingulate Cortex Area p24ab | 11.9 | 3.74 | -2, 36, 10 |
|  | 14.8 | 3.8 | R | Anterior Cingulate Cortex Area p24ab | 7.4 | 4.29 | 4, 32, 16 |
|  |  |  | L | Anterior Cingulate Cortex |  | 4.98 | -2, 28, 24 |
|  |  |  | L | Anterior Cingulate Cortex |  | 4.97 | -2, 24, 20 |
|  |  |  | L | Anterior Cingulate Cortex |  | 4.81 | -2, 20, 22 |
|  |  |  | R | Anterior Cingulate Cortex |  | 4.68 | 4, 24, 22 |
|  |  |  | L | Medial Cingulate Cortex |  | 4.42 | -6, 24, 32 |
| 266<br>ROI 2: dorsal striatum | 65.5 | 24.6 | R | Putamen (medial) | 10.6 | 4.79 | 20, 6, -4 |
|  | 25 | 9.4 | R | Putamen (bridges) | 8.5 |  |  |
|  | 21.6 | 8.1 | R | Caudate (bridges) | 6.8 | 4.41 | 18, 16, 2 |
|  | 19.6 | 7.4 | R | insula, Area Id1 | 10.9 | 4.13 | 38, -10, -10 |
|  | 18 | 6.8 | R | Putamen (lateral) | 6.4 |  |  |
|  | 12.5 | 4.7 | R | Globus Pallidus (extern) | 6.9 |  |  |
|  | 12.1 | 4.6 | R | Amygdala-striatal junction | 17.9 |  |  |
|  | 10.1 | 3.8 | R | insula, Area Id2 | 10.9 |  |  |
|  | 8.3 | 3.1 | R | Putamen (ventral) | 15.2 |  |  |
|  | 1.4 | 0.5 | R | Caudate (medial) | 0.5 |  |  |
|  | 1.3 | 0.5 | R | insula, Area Id3 | 0.5 |  |  |
|  | 0.9 | 0.3 | R | BF (Ch 4) | 2.5 |  |  |
|  | 0.6 | 0.2 | R | Globus Pallidus (ventral) | 0.7 |  |  |
|  | 0.4 | 0.1 | R | Amygdala (CM) | 1.1 |  |  |
|  |  |  | R | Putamen |  | 4.73 | 26, 4, -10 |
| 200<br>*ROI 3: TPJ* | 85.1 | 42.6 | R | Superior Temporal Gyrus<br>Area PGa (IPL) | 11.5 | 5.22 | 60, -52, 20 |
|  | 83 | 41.5 | R | Superior Temporal Gyrus<br>Area PF (IPL) | 12.2 | 4.63 | 64, -38, 24 |
|  | 25.3 | 12.6 | R | Area PFm (IPL) | 3.6 |  |  |
|  | 3.8 | 1.9 | R | Area PFcm (IPL) | 1.2 |  |  |
|  | 0.3 | 0.1 | R | Area PGp (IPL) | 0 |  |  |
| 180<br>*ROI 4: insula* | 14.3 | 7.9 | R | IFG (p. Opercularis) Area 45 | 1.3 | 4.25 | 54, 18, 8 |
|  | 11.6 | 6.5 | R | Area 44 | 1.9 |  |  |
|  | 2 | 1.1 | R | Putamen (medial) | 0.3 |  |  |
|  | 1.3 | 0.7 | R | Putamen (bridges) | 0.4 |  |  |
|  |  |  | R | Inferior Frontal Gyrus<br>(p. Triangularis) |  | 4.69 | 46, 28, 4 |
|  |  |  | R | Inferior Frontal Gyrus<br>(p. Opercularis) |  | 4.59 | 44, 16, 12 |
|  |  |  | R | Anterior insula Lobe |  | 3.83 | 36, 22, 4 |
| 165<br>*ROI 5: MTG* | 91 | 55.2 | R | Middle Temporal Gyrus<br>Area TE 4 | 20.6 | 5.46 | 52, -22, -8 |
|  | 37.5 | 22.7 | R | Area TE 5 | 5.3 |  |  |
|  |  |  | R | Middle Temporal Gyrus |  | 3.86 | 52, -36, -6 |

Only clusters surviving a 5% FWE correction at the cluster size are reported ( $t=3.37$ , $p < .001$ , cluster size 165). Brain regions are referred to as by the nomenclature of the Anatomy Toolbox (Eickhoff et al., 2005). Columns refer to the size in voxels of each cluster (voxels refer to the resampled 2x2x2mm voxels), the number of voxels of that cluster within a specific cytoarchitectonic region, the percentage of voxels in that region, hemisphere, cytoarchitectonic region (if available) or microanatomical region, percentage of cytoarchitectonic region falling within the cluster, peak t-value within a particular region and MNI coordinates of the peak, respectively. ROI 1 was further referred to as 'ACC'. ROI 2 was further referred to as dorsal striatum because over 50% of it fell
within the caudate/putamen. ROI 3 was further referred to as 'TPJ'. ROI 4 was further referred to as 'insula'. ROI 5 was further referred to as 'MTG'.

## RESULTS

### Behavioural results

#### Number of shocks delivered

In the coerced condition, participants were ordered to inflict 30/60 shocks, randomly. Two participants never followed the orders of the experimenter, one by inflicting 60/60 shocks, one by inflicting 0/60 shocks. They were fully removed from any further analyses. Three participants voluntarily disobeyed the orders to send a shock to the ‘victim’ on a few trials (respectively, 6 out of 30 shock trials, 5 out of 30 shock trials and 2 out of 30 shock trials). Those trials were removed from the analysis. In the free condition, participants were told that they were entirely free to decide to deliver a shock or not to the ‘victim’ on each of the 60 free trials. On average, agents administered 23.03/60 (SD=13.34, minimum: 1/60, maximum: 57/60) shocks, which was significantly lower than 30 (i.e. the number of shocks in the coerced condition, t_(36)_=−3.177, *p*=.003, Cohen’s d=−.522, BF_10_=11.70). There was no evidence for a difference across genders (24.60 for females and 19.75 for males, t_(35)_=1.036, *p*>.3, BF_10_=.504). Given the role reversal procedure in the present study, we assessed to what extent being first in the role of the ‘victim’ influences the subsequent choice to administer a shock to the new victim, previously agent. Previous studies indeed showed that participants who were victims first tend to administer more shocks than what they received when turning agent (Caspar et al., 2016; Caspar et al., 2017). A Pearson correlation between the number of shocks that participants received when they were victim first and the number of shocks that they gave when they became agents further showed a tendency to adapt the number of shocks they deliver to the number of shocks they received (r=.475, *p*_*1tailed*_ = .031, one-tailed, see **Fig. 3**). The Bayesian version of the same correlation also provides evidence for a positive correlation (BF_+0_=2.925). However, inverting the roles did not change the behaviour of participants, since an independent sample t-test revealed that the number of shocks sent by the participants did not differ according to their role order (victim first, agent first), t_(35)_=.802, p > .4. The Bayesian version of this test tends to support the absence of a difference (BF_10_=0.412). Those results thus show that the decision of participants to send shocks or not to the ‘victim’ is only slightly influenced by the role order during the task.

**Fig. 3.**
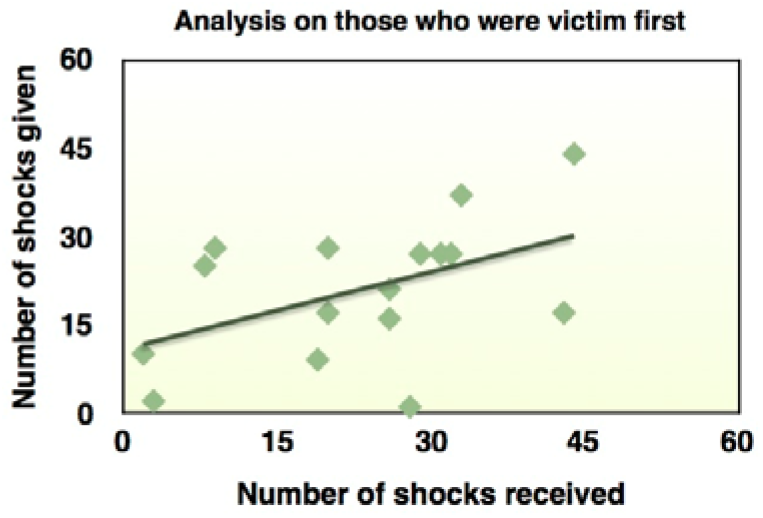
Pearson correlation between the number of shocks received for those who were victims first (n=16) and the number of shocks that they subsequently delivered when they turned agent.

#### Pain scale

A repeated-measures ANOVA with Condition (Free, Coerced) as within subject factor and Role-Order (Agent first, Victim first) as between-subject factors was used to analyse the pain ratings participants gave on 6 trials in each block. The scale ranged from ‘not painful at all’ (corresponding to ‘0’) to ‘very painful’ (corresponding to ‘1,000’). We observed a main effect of Condition (F_(1,29)_=9.921, *p* = .004, η^2^**_*partial*_** = .255). The Bayesian version of the same ANOVA resulted in a BF_incl_=5.580, confirming that the data is over 5 times more likely under models including Condition compared to those not including it, confirming what was found in the frequentist approach: the data provides evidence that pain values reported in the Free condition (501‰, CI_95_=425-577‰) exceeded those in the Coerced condition (457‰, CI_95_=382-532‰) (**Fig. 4A),** even if they knew that the shock intensity delivered to the ‘victim’ was the same throughout the entire experiment. We also observed an interaction between Condition and Role-Order (F_(1,29)_=6.303, *p* = .018, η^2^**_*partial*_** = .179). The Bayesian result also supports evidence for this interaction (BF_incl_=3.747). Independent sample t-tests indicated that the difference of pain ratings between the free and the coerced conditions was higher for those who were victim first (79‰, CI_95_=23-135‰) than for those who were agent first (9‰, CI_95_=−20 - 38‰, t_(29)_=−2.511, *p* = .018, BF_10_=3.415). The main effect of Role-Order resulted in a F_(1,29)_=0.004, *p* > .9 and a BF_incl_=1.484, suggesting that the effect size may be too small for our sample size and more data would likely be needed to make a clear statement about the absence or presence of a main effect of Role-Order. Given that participants delivered less shocks in the free condition (23.3/60) than in the coerced condition (30/60), reduced pain ratings in the coerced condition could be explained by a greater habituation to the pain response of the ‘victim’ in that condition in comparison with the free condition. Given that the number of shocks administered strongly varied between agents in the free condition, we performed the same repeated-measures ANOVA with the number of shocks freely delivered as a covariate. We observed that the pattern of results remained unchanged (see ***Supplementary Material S3***), thus confirming that differences in the pain ratings cannot be fully explained by a greater habituation to the pain response of the victim in the Coerced condition in comparison with the Free condition. Those results thus indicated that agents had a reduced perception of the pain of the ‘victim’ in the coerced condition compared to the free condition.

**Fig. 4.**
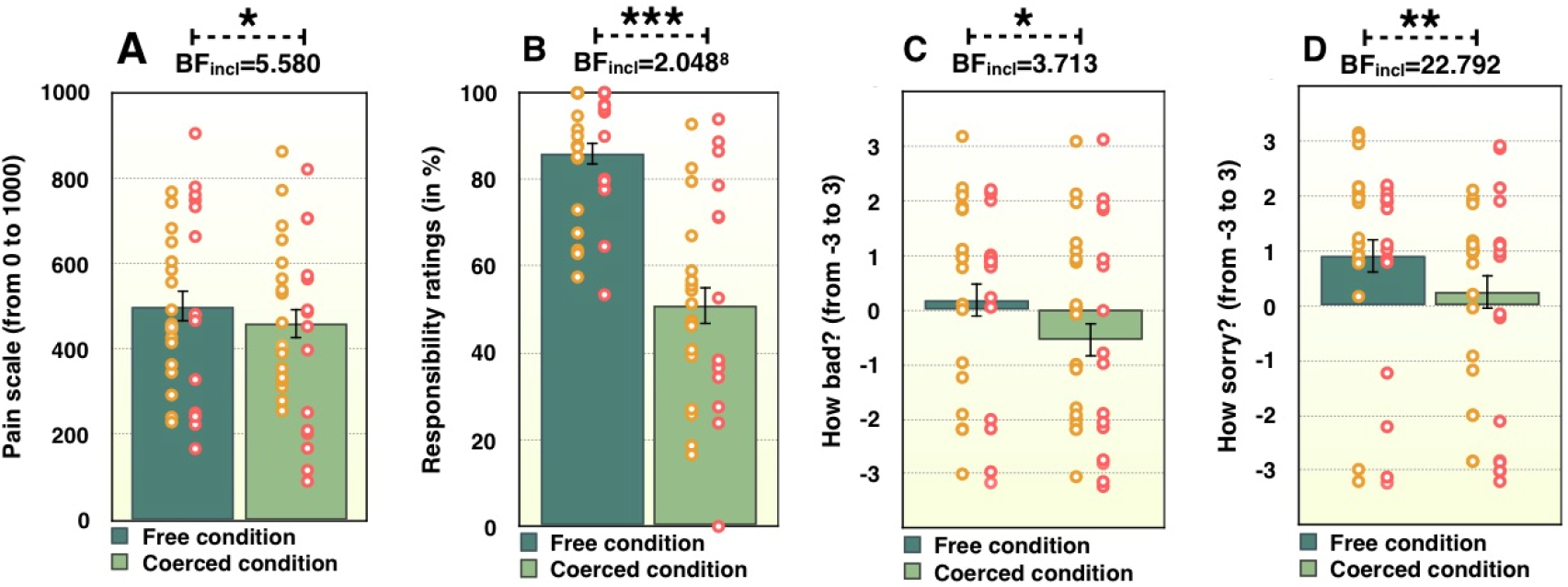
Behavioural results. Dark green columns represent the free condition and light green columns represent the coerced condition. A) Participants reported that the shocks administered to the ‘victim’ were more painful when they freely administered that shock than when they received the order to administer the same shock. B) Participants reported that they felt more responsible in the free than in the coerced condition. C) Participants reported that they felt worse administering a shock in exchange for money in the free condition than in the coerced condition. D) Participants reported that they felt sorrier to administer a shock in the free condition than in the coerced condition. All tests were two-tailed. Errors bars represent standard errors. *** indicates a p value <= .001. ** represents a p value between .001 and .01. * represents a p value between .01 and .05. BF refers to BF_incl_ in a Bayesian ANOVA. Colored dots represent individual participant data for agents first (orange) and victims first (red). For graphs C and D, a random jittering between −.25 and .25 has been added to each data point in order to avoid superposition.

#### Responsibility ratings

At the end of each of the four experimental runs, agents had to report how responsible they felt for the outcome of their actions. We conducted a repeated-measures ANOVA with Condition (Free, Coerced) as within subject factor and Role-Order (Agent first vs Victim first) as between-subject factor on the responsibility ratings. The main effect of Condition was significant, and Bayesian analyses strongly support the difference between the Free and Coerced condition (F_(1,31)_=59.696, *p* < .001, η^2^**_*partial*_** = .658; BF_incl_=2.048×10^8^). Participants reported a higher responsibility rating in the Free condition (86%, CI_95_=81-91%) than in the coerced condition (51%, CI_95_=42-60%) (**Fig. 4B)**. The main effect of Role-Order was not significant (F_(1,31)_=0.104, *p* > .7) and the Bayesian analysis provides evidence for the absence of a difference in the order the task was performed (BF_incl_=0.277). The interaction was not significant (F_(1,31)_=0.003, *p* > .9) and a Bayesian analysis provides evidence for the absence of an interaction (BF_incl_=0.301).

#### How bad and how sorry

At the end of the experiment, participants were asked to indicated how bad and sorry they felt separately for the free and the coerced condition when they administered shocks to the ‘victim’ on a scale ranging from ‘*Not sorry/bad at all*’ (−3) to ‘*Very sorry/bad*’ (3) (see ***Supplementary Material S2***). A repeated-measures ANOVA with Condition (Free, Coerced) as within subject factor and Role-Order (Agent first, Victim first) as between-subject factor on badness ratings was performed using both frequentist and Bayesian approaches. Results showed a main effect of Condition (F_(1,35)_=7.276, *p* = .011, η^2^**_*partial*_** =.172, BF_incl_=3.713). Participants reported that they felt worse in the free (.332, CI_95_=−.242 – 0.906) than in the coerced condition (−.448, CI_95_=−1.078 – .183) (**Fig. 4C)**. The main effect of Role-Order resulted in a F_(1,35)_=0.249, *p* > .6, BF_incl_=0.348 suggesting some evidence in support of the null hypothesis that role-order did not influence this rating. The interaction results also support the null hypothesis, as they resulted in not significant results with the frequentist approach (F_(1,35)_=0.011, *p* > .9) and a BF_incl_=0.312. The same ANOVAs on how sorry the agents felt again showed a main effect of Condition (F_(1,35)_=12.033, *p* < .001, η^2^_***partial***_ = .256, BF_incl_=22.792). Participants reported feeling sorrier in the free (.955, CI_95_=.375 – 1.536) than in the coerced condition (.025, CI_95_=(−).624 – .675) (**Fig. 4D)**. The main effect of Role-Order (F_(1,35)_=0.994, *p* > .3, BF_incl_=0.516) and the interaction (F_(1,35)_=0.192, *p* > .6, BF_incl_=0.510) were somewhat inconclusive, but did not provide evidence for an influence of Role-Order. The results thus indicated that participants felt worse and sorrier for the shocks sent to the ‘victim’ in the free condition compared to the coerced condition.

### MRI results

The key hypothesis of the present study was to observe reduced activity in regions associated with pain, including the insula and the ACC, while witnessing the pain of the victim in the coerced condition in comparison with the free condition. To identify the brain regions that respond more to the victim’s pain when people are free to decide to inflict pain compared to when they receive the order to inflict pain, we used the [FreeShock-FreeNoShock - (CoercedShock-CoercedNoShock)] contrast. Five participants only administered a very small number of shocks (between 1 and 5) which would not allow a reliable estimate of the brain activity in the FreeShock condition, and hence in our contrast of interest. These participants were thus not taken into account in any neuroimaging results (N=5, 3 were agents first). After excluding those participants, all remaining participants provided at least 8 shock trials, which appears to be enough to reveal regions with reduced shock-triggered brain activation when people obeyed orders compared to when they could freely decide which action to execute (t=3.37, *p* < .001, cFWE, i.e. cluster size threshold determined by 5% family wise error=165). These regions include the anterior cingulate cortex, putamen and caudate, the MTG, inferior parietal lobule, TPJ (overlapping with the TPJ in the meta-analysis of Mar, 2011), inferior frontal gyrus, insula (including anterior and dysgranular sections), and middle temporal gyrus (See **Table 1** and **Fig. 5**). Decoding the unthresholded map using Neurosynth revealed that the two functional terms most associated with this pattern of activation are pain and painful (correlated at r=0.211 and r=0.174, respectively, see https://neurosynth.org/decode/?neurovault=OEFCMEPO-390176). The reverse contrast [CoercedShock-CoercedNoShock - (FreeShock-FreeNoShock)] was not significant (for a t=3.37 and *p* < .001). Those results thus supported the main hypothesis, by showing reduced activity in brain regions associated with the processing of pain in the coerced condition compared to the free condition. To shed further light on the properties of the clusters identified in this contrast, we used the clusters as regions of interest (ROIs) to perform a number of follow up analyses. We conducted independent sample t-tests with order of the role as the independent factor and activity in our contrast of interest as the dependent variable in each of the ROIs. To correct for multiple comparisons for frequentist statistics, we applied a False Discovery Rate (FDR) approach with the Benjamini and Hochberg method (Benjamini & Hichberg, 1995) to each p-value with the frequentist approach. Results were inconclusive in all the ROIs (all *p*s_*FDR*_ > .1, all BFs_10_ > .41 & < 1.200). We also found no effect of gender in our contrast of interest in these ROIs (all *p*s > .1 and all BFs_10_<0.77&>.37)

**Fig. 5.**
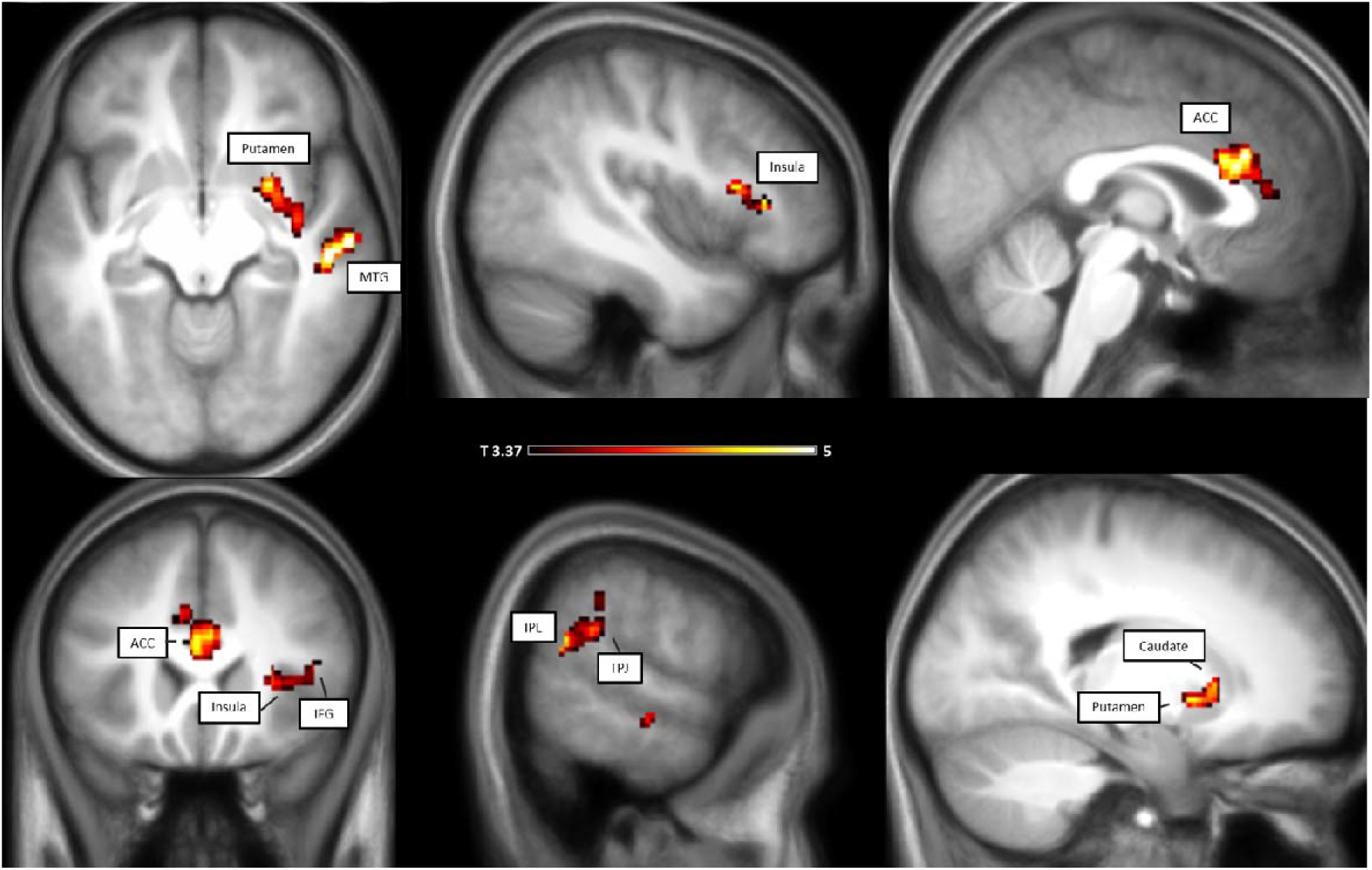
MRI results. [FreeShock-FreeNoShock - (CoercedShock-CoercedNoShock)] contrast. Peak coordinates can be seen in Table 1. Results are corrected using 5% FWE at cluster level (165 voxels), t=3.37, *p* < .001 at voxel level)

To further test our hypothesis that coercion leads to a reduced pain representation of the victim compared to the free condition, and to overcome the limitations of voxel-wise reverse inference, we used the vicarious pain signature identified by Krishnan et al., (2016). Krishnan and co-workers presented participants with images that suggested three levels of pain in other individuals, and identified a pattern of brain activity that predicts the level of pain (low, medium, high) perceived in the pictures based on the whole brain activity of the participant, even after exclusion of the occipital cortex. This vicarious pain signature (VPS) has been shown to successfully decode the level of pain attributed to characters in photographs across participants and studies (Krishnan et al., 2016). For the 32 participants that had sufficient trials in the free condition, we thus took the contrast [FreeShock - FreeNoShock] and the contrast [CoercedShock - CoercedNoShock] of each participant, and dot-multiplied each of them with the VPS, as described in Krishnan et al. (2016). We expected this decoded vicarious pain intensity to be lower in the coerced compared to the free condition, and therefore used a one-tailed t-test to test this prediction. We found that indeed, the VPS for the coerced Shock-NoShock (mean±sem, 15±7) was significantly reduced compared to the VPS for the free choice condition (36±9, t_(31)_=1.97, p_1tailed_=0.029, BF_+0_=1.998). We followed the same procedure using the Neurologic Pain Signature (NPS, Wager et al., 2013) confirming lower pain intensity in the coerced Shock-NoShock (mean±sem, −10.781±5.920) compared to the free condition (7.581±7.347, t_(31)_=2.044, *p*_1tailed_=0.025, BF_+0_=2.283). It should be noted that these effects are significant but modest in size, as evidenced by the BF~2. These results dovetail with the neurosynth decoding in associating the pattern of activity with painfulness.

Since we observed that participants reported that they felt less sorry and less bad for the shocks sent to the ‘victim’ in the coerced condition compared to the free condition, we further assessed whether coercion reduced neurocognitive processes associated with guilt. To do so, we applied the same procedure to a multivariate brain pattern for interpersonal guilt (Yu et al., 2020). A paired sample t-test comparing the guilt signature under coercion (−0.66±0.937) with that under free conditions (3.02±1.1) provides strong evidence for a reduction under coercion (t_(31)_=3.00, *p*_1tailed_=0.003, BF_+0_=15.3).

Participants sent on average less shocks in the free condition than in the coerced condition, which could thus have led to a higher repetition suppression to viewing shocks in the coerced condition. We thus performed additional analyses examining whether we had observed significant repetition suppression in our conditions. For that aim we compared brain activations when participants witnessed a shock that was preceded by another shock or by no shock, separately for the free and the coerced condition. This analysis did not show significant results in either condition at *p*<0.001 (cluster FWE corrected), thus implying that repetition suppression was not a significant factor in our analysis and that results are unlikely to be explained by differential repetition suppression.

### Region of Interest (ROI) approach based on the interaction

#### Decision-making in the free condition

If people would be more willing to perform moral transgressions under orders because orders reduce activations in regions associated with empathy that inhibit aggression, we would expect that activation level in those brain regions showing reduced activity under orders should be related to the decision-making to shock or not to shock. Given our low disobedience rate, we cannot test this notion in the coerced condition. Instead, we therefore tested whether individual differences in decision-making in the free condition (quantified as the number of shocks delivered to the ‘victim’) was related to individual differences in activity in empathy related brain regions showing reduced activity following orders (i.e. Table 1). Specifically, we thus estimated the contrast [FreeShock - FreeNoShock] from the average activity in each of the clusters of the contrast [FreeShock-FreeNoShock - (CoercedShock-CoercedNoShock)] (which we will now call ROIs) and used this parameter estimates in a correlation analysis with the number of shocks given.

We had prior hypotheses that based on findings relating brain activity in these regions to empathy and empathy to prosociality, individuals that show higher activity when witnessing a shock in these regions would decide to give fewer shocks (see Hein et al., 2010 and Gallo et al., 2018). We thus performed one-tailed correlations between the number of shocks that participants freely delivered and activity in the brain regions that we identified in the [FreeShock-FreeNoShock] contrast, again using both frequentist and Bayesian approaches. To correct for multiple comparisons for frequentist statistics, we applied a False Discovery Rate (FDR) approach with the Benjamini and Hochberg method (Benjamini & Hichberg, 1995) to each p-value with the frequentist approach. We performed separate analyses on those who were agent first and those who were agent second, since the choice to send or not a shock to the ‘victim’ for the later can be influenced by the number of shocks that they received as victim (see **Fig. 3**).

For those who were agent first, we observed that the number of shocks that they freely decided to administer was correlated with the activity in the dorsal striatum (i.e., ROI 2) (r=−.580, *p*_*FDR1tailed*_= .015), in the MTG (i.e., ROI 5) (r=−.579, *p_FDR1tailed_* = .015) and in the TPJ (i.e., ROI 3) (r=−.463, *p*_*FDR1tailed*_= .043). A Bayesian analysis also provides evidence for a negative correlation in the dorsal striatum (BF_−0_=11.062), the MTG (BF_−0_=10.882) and the TPJ (BF_−0_=3.217). There was however no conclusive evidence that activity was correlated with shocks with the ACC (ROI 1, *p*_*FDR1tailed*_> .1, BF_−0_=0.665) and the insula (ROI 4, *p*_*FDR1tailed*_ > .1, BF_−0_=0.843), see **Fig. 6A**. Supplementary Table S1 (A and B) reports clusters in which at the contrast [FreeShock-FreeNoShock] correlates with the number of free shocks delivered, including only agents first. Figure S1 displays the uncorrected results.

**Fig. 6.**
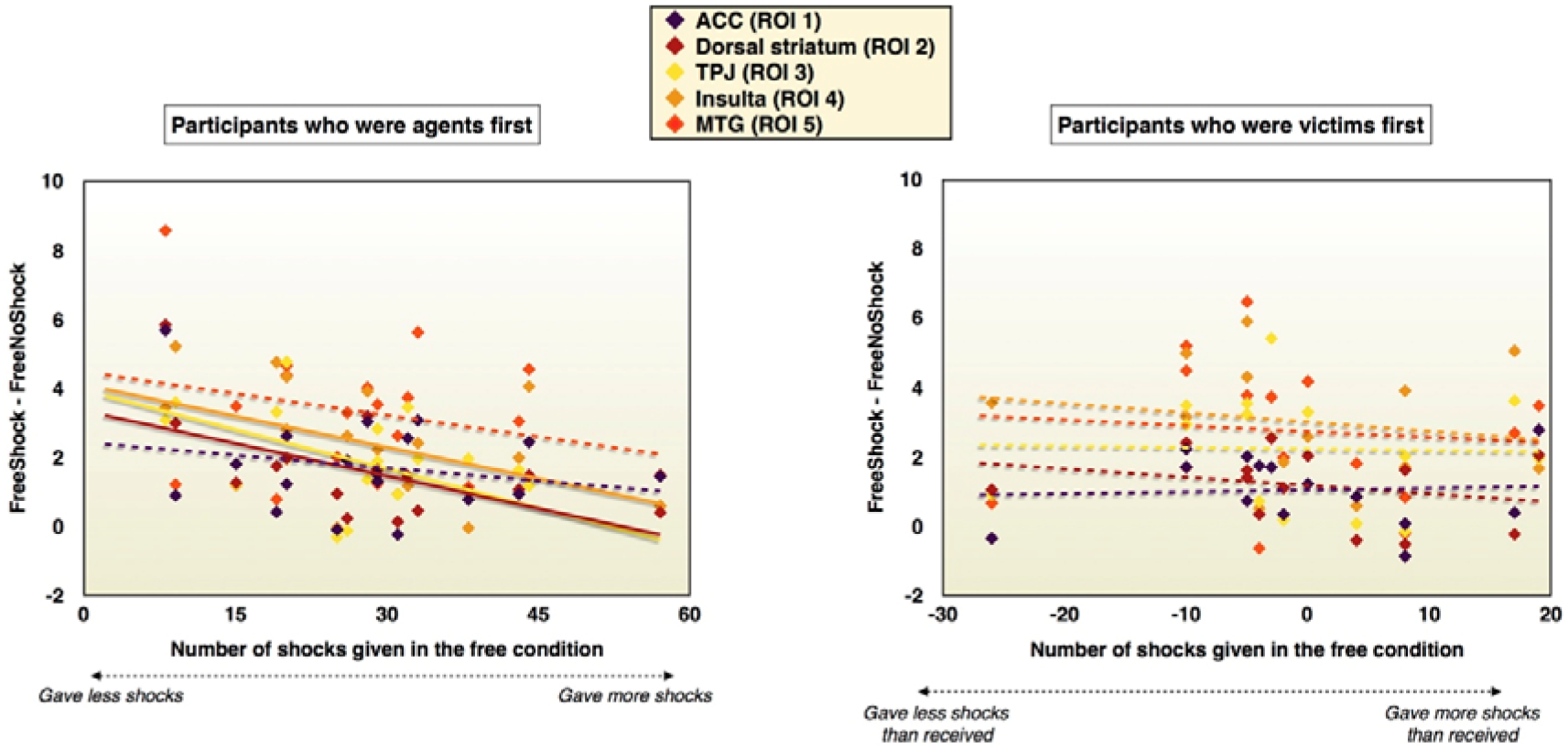
A) Relation between the number of shocks delivered in the free condition (x-axis) and ROIs for participants who were agent first (y-axis) in the contrast [FreeShock-FreeNoShock]. B) Relation between the number of shocks delivered in the free condition (x-axis) and ROIs for participants who were agent second (y-axis) in the contrast [FreeShock-FreeNoShock]. Full lines represent BF_−0_> 3, one-tailed.

For those who were agent second, we know that the number of shocks administered depends on the number of shocks received. To test whether there was additional information in brain activity, we subtracted the number of shocks received (as a victim) from the number of shocks delivered in the free condition (i.e., shocks given – shocks received), and correlated this excess with the brain activity in the regions of interest. There was no conclusive evidence that activity was correlated with excess shocks delivered in the dorsal striatum (*p* > .2, BF_−0_=0.791), the insula (*p* > .3, BF_−0_=0.424), the MTG (*p* > .4, BF_−0_=0.354) or the TPJ (*p* > .2, BF_−0_=0.569) (**Fig. 6B)**. For ACC the BF_−0_=0.285 suggests that data support the null hypothesis of a lack of correlation. Supplementary Table S2 (A and B) reports clusters in which at the contrast [FreeShock-FreeNoShock] correlates with the number of free shocks delivered, including only agents that were victim first. Figure S2 displays the uncorrected results.

#### Individual differences in empathy and the main effect of observing the victim’s pain

We also investigated to what extent the main effect of observing another individual’s pain (i.e., [FreeShock+CoercedShock - (FreeNoShock+CoercedNoShock)] contrast), irrespective of the experimental condition, was associated with individual differences in self-reported empathy. For this purpose we performed a multiple linear regression explaining the shock observation activity in the ROIs - measured through the [FreeShock+CoercedShock - (FreeNoShock+CoercedNoShock)] contrast - as a linear combination of an intercept and the score on the IRI for the empathy related subscales PT, EC and PD. (see Table 2). Analyses were performed using Bayesian Linear Regressions in JASP for the BF_incl_ and a frequentist Linear Regression in JASP for the partial correlation (*r*_p_) and the p value. Default priors implemented in JASP were used throughout. Because we include all three subscales in the multiple regression, we do not need to correct for the number of subscales. We did not correct for the number of ROIs, because such corrections are unusual when using Bayes factors, but this should be taken into account when considering the strength of the trait-brain-activity association. We tested normality for all subscales and ROI activity using the Shapiro-Wilk, and most did not violate normality (all *p*>0.14), except PD (*p*=.034) and the dorsal striatum (i.e., ROI 2, *p*=.045) and all Q-Q residual plots seemed reasonable. Considering the conceptual advantages of multiple linear regressions over pair-wise non-parametric association we therefore present the results of the linear regressions despite these deviations from normality, but results should again be interpreted with care. The Bayesian analyses revealed that activity in the ACC was associated with Empathic concern (i.e., EC), see **Fig.7**. It also provided evidence that TPJ activity was independent of EC and Personal Distress (i.e., PD). All other associations were inconclusive suggesting that our study was underpowered to provide evidence for or against those associations.

**Fig. 7.**
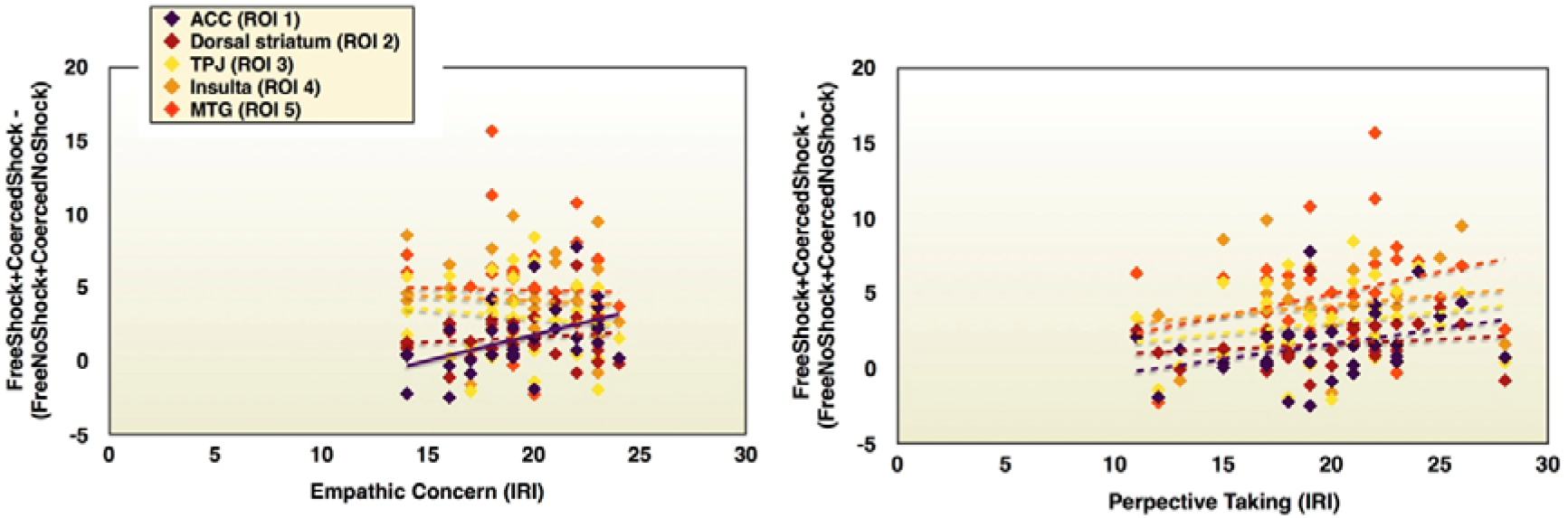
A) Graphical representation of the association between EC and ROIs (ACC: *p*=.016, BF_incl_=4.001) B) Graphical representation of the association between PT and ROIs. Full lines represent significant *p* values (*p* < .05) and BF_incl_ > 3, two-tailed.

**Table 2:** The relationship between shock activity in each ROI and trait measures of empathy. Results of the multiple linear regression explaining the shock observation activity ([FreeShock+CoercedShock - (FreeNoShock+CoercedNoShock)] contrast) in the ROIs as a linear combination of an intercept and the score on the IRI for the empathy related subscales PT, EC and PD. BF_incl_ refers to the Bayes factor for including a given subscale, *r*_p_ the partial correlation between the subscale and the activity in the ROI, and *p* the p value for including the subscale according to the linear regression. Scales for which the Bayes factor provides at least moderate evidence for inclusion (BF_incl_>3) are marked in green, scales for which the Bayes factor provides evidence for exclusion (BF_incl_<1/3) are marked in red.

| ROI | PT | EC | PD |
| --- | --- | --- | --- |
| **1 - ACC** | BF<sub>incl</sub>=1.368<br>$r_p$=0.237<br>$p$=0.206 | BF<sub>incl</sub>=4.001<br>$r_p$=0.436<br>$p$=0.016 | BF<sub>incl</sub>=0.953<br>$r_p$=-0.173<br>$p$=0.359 |
| **2 - Dorsal striatum** | BF<sub>incl</sub>=0.466<br>$r_p$=0.031<br>$p$=0.871 | BF<sub>incl</sub>=0.534<br>$r_p$=0.190<br>$p$=0.314 | BF<sub>incl</sub>=1.070<br>$r_p$=-0.345<br>$p$=0.062 |
| **3 - TPJ** | BF<sub>incl</sub>=0.348<br>$r_p$=0.184<br>$p$=0.331 | BF<sub>incl</sub>=0.284<br>$r_p$=-0.095<br>$p$=0.617 | BF<sub>incl</sub>=0.279<br>$r_p$=-0.005<br>$p$=0.978 |
| **4 - Insula** | BF<sub>incl</sub>=0.839<br>$r_p$=0.353<br>$p$=0.056 | BF<sub>incl</sub>=0.395<br>$r_p$=-0.122<br>$p$=0.522 | BF<sub>incl</sub>=0.406<br>$r_p$=0.147<br>$p$=0.439 |
| **5 - MTG** | BF<sub>incl</sub>=0.434<br>$r_p$=0.234<br>$p$=0.214 | BF<sub>incl</sub>=0.359<br>$r_p$=-0.175<br>$p$=0.354 | BF<sub>incl</sub>=0.313<br>$r_p$=0.017<br>$p$=0.929 |

### Increased activity during the Free decision-making phase

Using the predictor starting with the order and ending with the button press, in an exploratory analysis we also analysed whether there were differences in activity during the decision-making phase of the experiment between Free and Coerced conditions. This predictor included the decision-phase of all trials, independently of whether participants later chose to give or not to give shocks. At p<0.001 5%FWE cluster corrected, we observed a cluster including the left superior medial and mid orbital gyri in the vmPFC that was more active while taking a decision in the Free compared to following orders in the Coerced condition (See **Table S3** and **Fig. S3**). This finding matches the fact that participants had to engage in valuational decision-making in the Free condition, but doing so in the Coerced condition was optional, and such decision-making has been repeatedly associated with the vmPFC, particularly when participants need to decide how much money to give up to benefit others (Hare, Camerer, Knoepfle, & Rangel, 2010; Hutcherson, Bushong, & Rangel, 2015; FeldmanHall, et al. 2015). No clusters survived the reverse contrast (i.e. more activity during Coerced than Free decisions).

## DISCUSSION

### Reduced vicarious activations in the coerced vs free condition

We tested the hypothesis that obeying the orders received from an authority would reduce the vicarious brain activation when witnessing the pain that one had delivered to a ‘victim’ compared to a free condition. MRI results confirmed this hypothesis: when participants witnessed a shock that they delivered after having received the order to do so, activity in brain regions including ACC, dorsal striatum (putamen and caudate), MTG, TPJ and insula/IFG was reduced in comparison with being free to decide to deliver that pain. Participants knew that the shock intensity was kept constant during the entire experiment, no matter the experimental conditions, yet activity in empathy-related brain regions was attenuated in the coerced condition. Considering the difficulty of unambiguously associating activity in specific brain regions with mental processes (Poldrack 2006), it is important to notice that using a vicarious pain signature developed by Krishnan et al. (2016) to decode perceived pain intensity from brain activity and using a neurological pain signature developed by Wager et al., 2013 to decode felt pain intensity lead to the same conclusion. Importantly, additional analyses confirmed that reduced activations in the coerced condition were not related to a repetition suppression phenomenon associated with viewing shocks more frequently in the coerced condition than in the free condition.

Participants also reported a reduced feeling of responsibility in the coerced condition, which may have contributed to the decreased processing of the consequences of the agent’s actions on the victim. This dovetails with previous studies that reached a similar conclusion (e.g. Cui et al., 2015; Koban et al., 2013; Lepron et al., 2015), but goes beyond by showing that even in the case of a pain that is fully caused by the participants own action, brain activity is altered by a lack of responsibility. For instance, in Lepron et al. (2015) the only statistical differences observed were between two conditions (‘observe’ vs ‘decide and execute’) that not only differed in terms of responsibility, but also in the sensorimotor information associated with performing an action, in the causality between one’s own actions and the outcome and in the degree of control associated with the action. In Cui et al. (2015), the painful outcome delivered to the recipient depended on the participant’s errors but also on the recipient’s errors. In our paradigm, participants were always the authors of the actions and these actions were always fully predictable, thus isolating the impact of one’s own perception of responsibility on the empathic response.

It may be argued that our results do not only reflect a reduced empathic response when people obey orders but a general effect of lack of freedom to make a decision. Such conditions were already introduced in a previous study (Lepron et al. 2015) in which participants also had to perform a condition in which they had to execute an instruction sent by the computer. Interestingly, the authors did not observe any differences in the neurophysiological empathic response between that condition and a condition in which participants could decide and execute the action. In our study, we did observe such differences at the neural level between the free and the coerced condition, suggesting that obeying the orders of an authority has a stronger influence on the empathic response towards others’ pain than simply following the instructions of a computer.

A study (Yu et al., 2014) showed that the activations in anterior middle cingulate cortex (aMCC) and anterior insula (AI) were higher when participants felt guiltier and were the sole responsible for the pain of others compared to when they bear less responsibility. In the present study, participants explicitly reported that they felt less sorry and less bad in the coerced condition compared to the free condition. The reduced activations observed in the ACC and insula could thus also reflect a reduced perception of guilt under coercion. This notion was confirmed by the fact that a multivoxel pattern that has been shown to decode interpersonal guilt from neural activity (Yu et al., 2020) was also reduced in our Coerced vs. Free condition.

### Sense of responsibility vs sense of agency

However, we could not disentangle if a reduced feeling of responsibility accounted for the entire effect or if this relationship could be mediated, at least in part, by a reduced sense of agency, since both are influenced by obedience to an authority (Caspar et al., 2016). The sense of agency is the feeling that we are the authors of our own actions and of their consequences (Gallagher, 2000). One measure that has been used in past literature to assess the implicit feeling of agency is based on time perception (Haggard et al., 2002) between an action and its outcome. Unfortunately, if the consequence of an action occurs more than 4s after the action, modulations of the sense of agency no longer lead to measurable changes in time perception (Humphrey & Buehner, 2009). In order to separate brain activity related to motor response from those related to processing the pain of the victim, comparatively long jittered action-outcome delay had to be used in our *f*MRI paradigm, precluding the use of time-estimates to probe the sense of agency in the present paradigm. Future studies could however aim to disentangling the effects of sense of agency and responsibility by using conditions in which brain activity is measured from individuals giving orders, which gives a similar level of responsibility but a reduced sense of agency (Caspar et al., 2018).

### Pain is perceived as less intense in coerced vs free conditions

Strikingly, even if participants were explicitly told that the shock intensity would remain the same throughout the entire experiment for the victim, they rated the shock intensity lower in the coerced condition than in the free condition, mimicking the decoded vicarious pain intensity. In a previous study, Akitsuki & Decety (2009) observed that participants reported higher pain ratings when they witnessed a painful stimulation that was caused on purpose by another individual than when that pain was triggered by accident by the person itself, suggesting that intentionality plays a role in the subjective evaluation of pain. However, in Akitsuki & Decety (2009) participants were not involved in what happened on those videos, they were simple witnesses. In addition, they were not explicitly told that the shock intensity would be the same during the experimental conditions. Here, our results would support the fact that obeying orders has such a strong influence on the perception of pain felt by others that it even impacts perceptual reports of observed shock intensity rather than only modulating how the observer feels about the pain of the other.

### Vicarious activations covary with the number of shocks given

Previous studies on prosocial behaviours observed that activity in the regions associated with the pain network, mainly the ACC and the AI, predicted helping behaviours (Hein et al., 2010; FeldmanHall et al., 2015). Animal studies have also revealed that deactivating the ACC increases the number of shocks a rat is willing to give to another rat (Hernandez-Lallement et al., 2020). Here, we predicted the same effect but did not find that the activity of these two brain regions was related to prosocial behaviours in our task (ACC - ROI 1 and insula - ROI 4). Rather, we observed that the activity in the dorsal striatum (ROI 2), in the MTG (ROI 5) and in the TPJ (ROI 3) was related to how prosocially participants acted in the free condition, by administering fewer shocks if activity in these regions to witnessing the victim receive a shock was higher.

We observed that the number of shocks that agents freely chose to deliver was associated with the activity in the TPJ. The TPJ is a brain region associated with emotional perspective taking and theory of mind (Ruby & Decety, 2004; Young et al., 2007) and has been linked to prosocial behaviours (FeldmanHall et al., 2015). It has been argued that highly aggressive participants show cognitive disengagement, associated with reduced mentalizing about the consequences of their actions on their co-participant (Zaki & Ochsner, 2012). Additionally, the striatum has been shown to respond to both monetary and social rewards (e.g. Izuma, Saito & Sadato, 2008; Knutson, et al. 2001; Saxe & Haushofer, 2008) and is linked to prosocial behaviours (e.g. Delgado et al., 2005; Kokal et al., 2011). We also observed that the more the middle temporal gyrus was activated, the less shocks participants freely delivered. Previous studies already observed that the middle temporal gyrus is involved when participants witness painful vs non-painful stimuli (e.g. Coll, Grégoire, Eugène & Jackson, 2017; Lang, Yu, et al. 2011), thus suggesting that a higher activity of the MTG is associated with a higher sensitivity to the pain of others. A review further proposed a model where higher activity in the MTG modulates aggressive behaviour (Potegal, 2012), which would be consistent with the present results.

However, since our sample was split in two; those who were agent first and those who were ‘victim’ first, the final sample size to perform correlations was quite low (i.e. N was between 16 and 21). Bayes Factors confirmed a lack of sensitivity for some of the correlations, probably due to this low sample size. Future studies evaluating similar research questions could thus integrate a higher sample size for testing the order of the role of participants across correlations.

With the role reversal in our task, we observed that participants who had been victims first tended to incorporate the number of shocks received in their decision-making, with a significant correlation between shocks received and shocks later given. This effect has already been observed in former studies based on a similar paradigm (Caspar, et al., 2016; 2017). One explanation for this effect is vindictiveness: that participants who have received many shocks deliver more shocks to punish the new victim. Another possibility is that participants who were victims first tended to administer the same amount of shocks or slightly more to the ‘victim’ when they turned to be agents because they perceived that it would be fairer to be paid equally. The present result could thus reflect vindictiveness, in the sense of a wish to punish, or it could reflect that participants show inequity aversion, and are thus keen to deliver a number of shocks that is ‘fair’ given those they have received. However, this ‘vindictive tendency’ was smaller in the present study than in former studies (e.g. Caspar et al., 2016), possibly because ‘victims’ were less able to “count” the number of shocks received in the free condition because of the split of conditions in different MRI runs and the longer timing between each event.

### Conclusion

Milgram’s famous experiments have shown widespread willingness to obey authority, even to the point of inflicting harm to an innocent volunteer (Milgram, 1963). The mechanisms underlying such behaviour, that is frequently justified by the fact that the person was simply obeying orders and thus did not feel responsible, are still largely unexplored despite their high social relevance. Here, we have shown that when people accept to comply with the orders of an authority, the neural response associated with the perception of pain felt by another individual is reduced in comparison with being free to choose which action to perform. Crucially, even subjective pain ratings indicated that the pain delivered to the other individual appeared less painful when people administered the shock according to the experimenter’s order than when they could freely decide. We also show that participants’ negative feelings and neural guilt signatures were reduced when they comply with orders. These results highlight how obeying an order relaxes our aversion against harming others, despite still being the author of the action that led to the pain.

## Acknowledgements

The research was funded by the European Union’s Horizon 2020 research and innovation programme under the Marie Skłodowska-Curie grant agreement Agent No 743685 to E.A.C., and from the Netherlands Organization for Scientific Research (VICI: 453-15-009 to C.K. and VIDI 452-14-015 to V.G.).

