## Supplementary for "Obeying orders reduces vicarious brain activation towards victims’ pain"

SUPPLEMENTARY MATERIAL S1

“ In this experiment, there will be two roles: One of ‘agent’ and one of ‘victim’. We will select randomly who starts with which role by picking a card in a box, but you can also tell us if you have a preference to start either by the role of the agent or by the role of the victim. Your roles will be switched at the middle of the experiment.”

“ The person in the role of the agent will be in the scanner and the person in the role of the victim will be located outside the scanner, in the control room. The agent will have two buttons: One button associated with SHOCK and one button associated with NO SHOCK. If the SHOCK button is pressed, the agent will send a mildly painful electric shock to the victim but will earn + €0.05. If the NO SHOCK button is pressed, there is no shock delivered to the victim and no additional money earned. In order to know which button is associated with shock or no shock, you will see two rectangles appearing on the screen on each trial: A green one labelled SHOCK and a red one labelled NO SHOCK. You will have to press the button corresponding to the outcome you want. The position of those buttons will vary randomly on a trial-basis but the outcome will always be fully congruent with what you see on the screen. As an agent, you will have two experimental conditions: One condition in which you can freely decide which button to press and another condition in which the experimenter will tell you which button you have to press. You will be able to hear the instructions of the experimenter through headphones that you will wear when you will be in the MRI screen. The instructions will be either “give a shock” or “don’t give a shock”. The experimenter will be with you in the scanner room to give her instructions. She will wear a microphone so you will still be able to hear her despite the noise of the MRI scanner. In the condition in which you can freely decide what to do, the experimenter will tell you, before each trial, another verbal instruction, which is “you can decide”. During that condition the experimenter will be located in the control room and will give the orders through the interphone.”

“Before we start the experiment, we will determine your own pain threshold. This experiment indeed involves sending real shocks to the other participant but those shocks will be calibrated. We will place two electrodes on your left hand and will connect them to a machine that is commonly used in physiotherapy to stimulate muscles or nerves. Here we will stimulate your muscles. We will increase the threshold step by step: At the beginning you won’t feel anything. Then, at some point you will be able to feel something very small, this is your detection threshold. From that threshold, we will continue to increase. The sensation will be weird and you will have muscle twitches after a certain threshold. Those twitches are normal since we stimulate a muscle. We will continue to increase until we reach your own pain threshold. We will ask you several questions to ensure that we have reached the pain threshold. When you will be in the role of the victim, this is the threshold that you will feel each time you receive a shock. This threshold will never increase or decrease during the experiment. There is absolutely no risk of burns or any other long-term undesired effects with this procedure.”

“When you are in the role of the agent, you will be ‘isolated’ in the scanner. Therefore, we have placed a camera above the hand of the victim that will display in real-time the hand of the victim inside the scanner on the screen that will be positioned in front of you. You will thus be able to see the outcome of your actions. It is very important that you always look at the hand of the victim when you perform the task. Therefore, you will see an arrow pointing to the top after you press the button in order to remind you to look at the hand of the victim. You will also see, on some trials, a pain scale appearing on the screen. That pain scale will range from ‘not painful at all’ to ‘very painful’. You will have a marker appearing at a random position on that scale and you will be able to move the position of the marker by using the buttons below your middle fingers for fast move or below your index fingers for slow move. Stop the marker at the position that corresponds to your estimation of the pain of the victim on the trial that just happened. If that scale appears after a no shock trial, then put the marker on ‘not painful at all’, since there was no shock’. If there was a shock delivered on the hand of the victim, rate what you think the pain of the victim was. As a reminder, the pain of the victim will always be the same. This question is asked to ensure that you paid attention to the hand of the victim during the task, as requested. We will start with a training session.”

“When you are in the role of the victim, you will be sitting at a table in the control room with your left hand connected to two electrodes. Please do not move your hand at all during the recording to avoid biasing what the agent sees during the recording of his/her brain activity.”

“You will first have 30 trials in one experimental condition followed by 30 trials in the other condition. Then, we will run the anatomical scan so you can rest a bit. After the anatomical scan, you will have again 30 trials in one experimental condition followed by 30 trials in the other experimental condition. You will thus have a total of 60 trials in each experimental condition. When the agent finishes the experiment, we will switch your roles”.

SUPPLEMENTARY MATERIAL S2

**1.**	**How bad did you feel when you delivered a shock in exchange for money?**

In the free condition

(-3) ------ (-2) ------ (-1) ------ (0) ------ (+1) ------ (+2) ------ (+3)

    	Not bad at all                                                           	 	Very Bad

In the coerced condition

(-3) ------ (-2) ------ (-1) ------ (0) ------ (+1) ------ (+2) ------ (+3)

    	Not bad at all                                                           	 	Very Bad

**2.**	**How sorry did you feel when you delivered shocks to the ‘victim’?**

In the free condition

(-3) ------ (-2) ------ (-1) ------ (0) ------ (+1) ------ (+2) ------ (+3)

    	Not sorry at all                                             	               	Very sorry

In the coerced condition

(-3) ------ (-2) ------ (-1) ------ (0) ------ (+1) ------ (+2) ------ (+3)

    	Not sorry at all                                             	               	Very sorry

**3.**	**Please describe in a few word how did you feel during the experiment**

**4.**	**Which role (agent or victim) did you prefer and why?**

**5.**	**As an agent, which condition did you prefer (free or coerced) and why?**

SUPPLEMENTARY MATERIAL S3

| **Effects** | **Frequentist Results** | **Bayesian Results (BF_incl_)** |
| --- | --- | --- |
| Condition | F_(1,28)_=5.120, *p* = .032 | 6.144 |
| Order of the Role | F_(1,28)_=.005, *p* > .9 | 1.542 |
| Shocks delivered | F_(1,28)_=.168, *p* > .6 | .692 |
| Condition * Order of the Role | F_(1,28)_=6.192, *p* = .019 | 4.023 |
| Condition * Shocks given | F_(1,28)_=.859, *p* > .3 | / |

SUPPLEMENTARY FIGURE S1

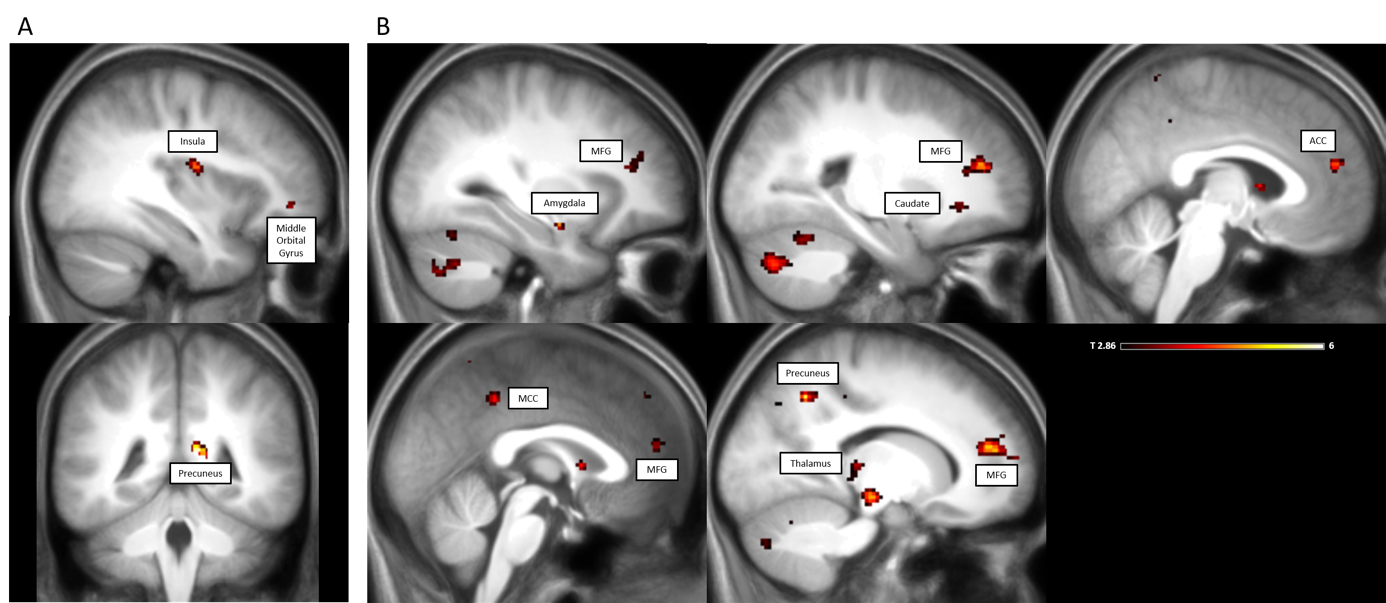

**Fig. S1.** **MRI results**. A) Regression analysis of contrast [FreeShock-FreeNoShock] using the number of shocks that participants who were agent first administered to the victim, uncorrected, t=2.86, *p* < .005. B) Regression analysis of the contrast [FreeShock-FreeNoShock] using the number of shocks that participants who were agent first administered to the victim, negative relationship, uncorrected, t=2.86, *p* < .005. Peak coordinates of the cluster surviving FWE correction at the cluster size can be seen in Table S1.

SUPPLEMENTARY TABLE S1

**TABLE S1.A - Agents first - Regression [FreeShock-FreeNoShock] - number of shocks delivered, *positive relationship, uncorrected***

| ***cluster size***  ***(N voxels)*** | **#Voxels in cyto** | **% Cluster** | **Hem** | **Cyto or Anatomical description** | **%**  **Area** | **cluster peak t-value** | **MNI coordinates (X,Y,Z)** |
| --- | --- | --- | --- | --- | --- | --- | --- |
| 60 |  |  | R | PPC |  | 6.32 | 10, -42, 14 |
|  |  |  | R | Precuneus |  | 4.18 | 18, -46, 12 |
| 44 | 24.3 | 55.1 | R | Area OP3 [VS]  Insula | 11.6 | 4.46 | 36, -10, 16 |
| 28 | 3.8 | 13.4 | R | Area Fp1 | 0.2 | 4.03 | 30, 46, 2 |
|  |  |  | R | Middle Orbital Gyrus |  | 3.66 | 38, 44, -6 |
| 22 | 8.1 | 36.9 | L | Area Ig2  Insula | 6 | 3.53 | -36, -16, 8 |
|  | 6.4 | 29 | L | Area OP3 [VS]  Insula | 4.5 | 3.00 | -38, -12, 14 |
|  | 1.3 | 5.7 | L | Area TE 1.0  Superior Temporal Gyrus | 1.0 | 3.15 | -44, -18, 4 |

This Table represents a summary of the main results. Only main peaks are reported. Clusters with voxels in cyto < 20 were not reported. Clusters with N/A areas only where not reported. Full results are displayed in a .txt file entitled TABLE S1.A on OSF.

**TABLE S1.Bi - Agents first - Regression [FreeShock-FreeNoShock] - number of shocks delivered, *negative relationship, uncorrected***

| ***cluster size***  ***(N voxels)*** | **#Voxels in cyto** | **% Cluster** | **Hem** | **Cyto or Anatomical description** | **%**  **Area** | **cluster peak t-value** | **MNI coordinates (X,Y,Z)** |
| --- | --- | --- | --- | --- | --- | --- | --- |
| 579 |  |  | L | Superior Medial Gyrus |  | 6.08 | -10, 46, 20 |
| 372 | 34.1 | 9.2 | L | Fusiform Gyrus | 4.1 | 4.86 | -40, -54, -22 |
|  | 163 | 43.8 | L | Cerebelum (VI) | 8.7 | 4.27 | -34, -66, -22 |
| 299 |  |  | L | Middle Temporal Gyrus |  | 5.64 | -52, -20, -12 |
| 293 | 136 | 46.4 | L | Cerebelum (Crus 2) | 8.3 | 4.27 | -22, -76, -38 |
| 243 | 143.8 | 59.2 | R | Cerebelum (Crus 1) | 4.4 | 4.49 | 22, -78, -30 |
| 218 |  |  | R | Middle Frontal Gyrus |  | 4.70 |  |
| 54 | 25.6 | 47.5 | L | Thalamus | 4.8 | 4.04 | -10, -30, 4 |
| 44 | 27 | 61.4 | R | Precuneus | 3.7 | 3.62 | 6, -52, 68 |
| 44 | 33.6 | 76.4 | L | Postcentral Gyrus | 6.4 | 4.11 | -42, -32, 50 |
| 44 |  |  | L | IFG |  | 4.53 | -36, 18, 32 |
| 41 | 28 | 68.3 | R | Cerebelum (Crus 1) | 1.6 | 4.27 | 42, -48, 30 |
| 36 | 23.3 | 64.6 | L | Postcentral Gyrus | 7.2 | 3.79 | -44, -18, 42 |
| 34 |  |  | L | Caudate Nucleus |  | 3.23 | -18, 24, -2 |
| 31 |  |  | L | MCC |  | 4.01 | 0, -42, 42 |
| 31 |  |  | L | Superior Medial Gyrus |  | 3.83 | -6, 42, 32 |
| 24 | 13.1 | 54.7 | L | Cerebelum (IX) | 14.6 | 3.65 | -4, -54, -38 |

This Table represents a summary of the main results. Only main peaks are reported. Clusters with voxels in cyto < 20 were not reported. Clusters with N/A areas only where not reported. Full results are displayed in a .txt file entitled TABLE S1.Bi on OSF.

**TABLE S1.Bii - Agents first - Regression [FreeShock-FreeNoShock] - number of shocks delivered, *negative relationship,  FWE corrected at cluster size***

| ***cluster size (N voxels)*** | **#Voxels in cyto** | **% Cluster** | **Hem** | **Cyto or Anatomical description** | **%**  **Area** | **cluster peak t-value** | **MNI coordinates (X,Y,Z)** |
| --- | --- | --- | --- | --- | --- | --- | --- |
| 128 |  |  | L | Superior Medial Gyrus |  | 6.08 | -10, 46, 20 |
| 121 |  |  | L | Middle Temporal Gyrus |  | 5.64 | -52, -20, -12 |
|  |  |  | L | Superior Temporal Gyrus |  | 4.50 | -52, -14, -6 |
|  |  |  | L | Middle Temporal Gyrus |  | 4.37 | -54, -16, -20 |
|  |  |  | L | Middle Temporal Gyrus |  | 4.35 | -54, -12, -18 |
|  |  |  | L | Middle Temporal Gyrus |  | 4.01 | -56, -10, -16 |

Only clusters surviving a 5% FWE correction at the cluster size are reported (t=3.58, *p* < .001, cluster size 121). Brain regions are referred to as by the nomenclature of the Anatomy Toolbox (Eickhoff et al., 2005). Columns refer to the size in voxels of each cluster (voxels refer to the resampled 2x2x2mm voxels), the number of voxels of that cluster within a specific cytoarchitectonic region, the percentage of voxels in that region, hemisphere, cytoarchitectonic region (if available) or microanatomical region, percentage of cytoarchitectonic region falling within the cluster, peak t-value within a particular region and MNI coordinates of the peak, respectively. The file is downloadable as a .txt file entitled ‘Table S1.Bii’ on OSF.

SUPPLEMENTARY FIGURE S2

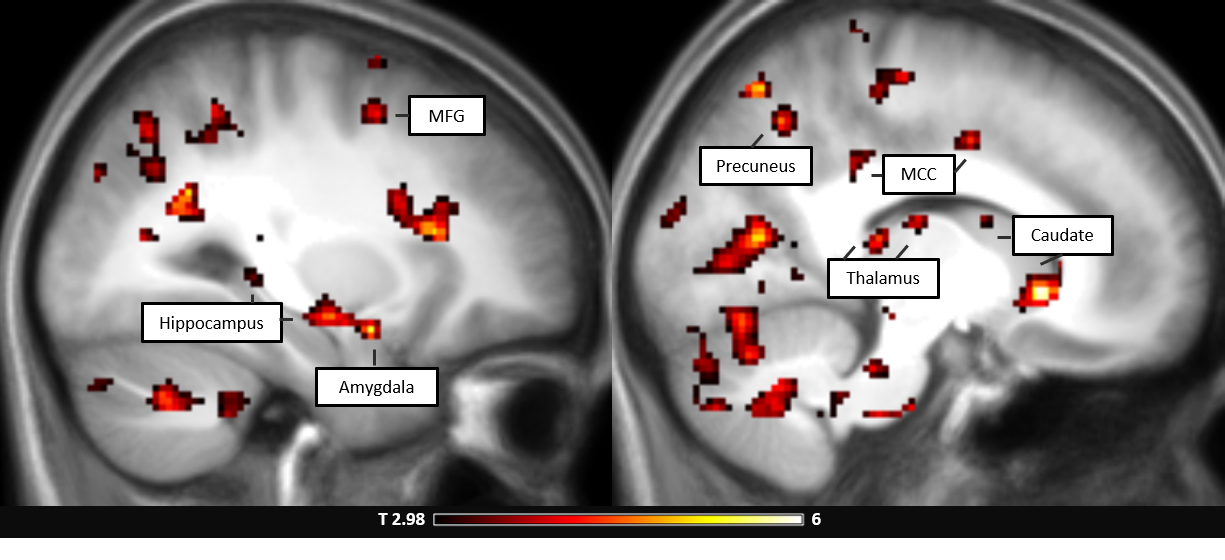

**Fig. S2.** **MRI results**. Regression analysis of contrast [FreeShock-FreeNoShock] using the numbers of shocks that participants who were victims first delivered to the victim minus the number of shocks received as a victim themselves, negative relationship, uncorrected, t=2.98, *p* < .005. Peak coordinates of the cluster surviving FWE correction at the cluster size can be seen in Table S2. The reverse relationship was not significant (for a t=2.98 and *p* < .005).

SUPPLEMENTARY TABLE S2

**TABLE S2.A - VICTIM FIRST - Regression  [FreeShocks-NoShock] - vindictiveness,  *negative relationship,  uncorrected***

| ***cluster size***  ***(N voxels)*** | **#Voxels in cyto** | **% Cluster** | **Hem** | **Cyto or Anatomical description** | **%**  **Area** | **cluster peak t-value** | **MNI coordinates (X,Y,Z)** |
| --- | --- | --- | --- | --- | --- | --- | --- |
| 15028 |  |  | L | Precuneus |  | 7.58 | -6, -62, 48 |
|  | 42 | 0.3 | L | Hippocampus | 19.1 | 7.27 | -36, -28, -12 |
|  | 175.4 | 1.2 | L | Thalamus | 55.1 | 6.52 | -18, -26, 4 |
| 564 | 16.6 | 2.9 | R | Caudate Nucleus | 7.7 | 6.11 | 10, 20, -4 |
| 503 |  |  | R | Caudate Nucleus |  | 5.55 | 16, 8, 22 |
| 431 | 265.6 | 61.6 | R | Cerebelum | 8.2 | 4.43 | 34, -62, -36 |
|  | 72.3 | 16.8 | R | Cerebelum VI | 4 | 3.45 | 30, -42, -36 |
| 367 | 35.11 | 9.6 | R | ParaHippocampal Gyrus | 9.3 | 4.48 | 24, -30, -12 |
| 332 | 31.9 | 9.6 | L | IFG | 4.6 | 4.31 | -48, 30, 24 |
| 222 | 26 | 11.7 | R | Superior Temporal Gyrus | 12.9 | 4.74 | 44, -32, 10 |
| 170 |  |  | L | Middle Frontal Gyrus |  | 4.03 | -42, 50, 12 |
| 141 | 10.3 | 7.3 | R | Fusiform Gyrus | 2.1 | 6.02 | 38, -30, -20 |
| 80 | 35.4 | 44.2 | R | Postcentral Gyrus | 5.4 | 4.43 | 42, -26, 40 |
| 74 | 24.9 | 33.6 | R | Middle Temporal Gyrus | 2.5 | 4.49 | 52, -60, 20 |
| 69 | 39.8 | 57.6 | R | Fusiform Gyrus | 12.2 | 4.14 | 38, -62, -14 |
| 65 |  |  | L | Middle Temporal Gyrus |  | 4.17 | -56, 4, 22 |
| 62 | 16.1 | 26 | L | Superior Temporal Gyrus | 4.3 | 4.10 | -50, -22, 10 |
| 58 |  |  | R | Middle Frontal Gyrus |  | 3.90 | 36, 36, 24 |
| 50 | 46.1 | 92.3 | R | Superior Temporal Gyrus | 11.8 | 4.22 | 48, -24, 16 |
| 45 | 15.4 | 34.2 | L | Postcentral Gyrus | 1.6 | 4.14 | -20, -28, 70 |
| 40 | 21.4 | 53.4 | L | Postcentral Gyrus | 3.8 | 4.03 | -46, -32, 60 |
| 39 | 21.1 | 54.2 | R | Postcentral Gyrus | 3 | 5.57 | 64, -10, 28 |
| 35 |  |  | L | IFG |  | 4.27 | -42, 50, -12 |
| 34 | 15.5 | 45.6 | L | Heschls Gyrus | 4.3 | 3.81 | -62, -10, 10 |
|  | 14.6 | 43 | L | Superior Temporal Gyrus | 3.9 | 3.98 | -60, -20, 12 |
| 25 |  |  | R | Middle Temporal Gyrus |  | 3.61 | 62, -46, -8 |
| 24 |  |  | R | MCC |  | 4.26 | 12, -2, 42 |
| 23 | 11.9 | 51.6 | L | SupraMarginal Gyrus | 2.1 | 4.14 | -62, -22, 38 |
| 23 | 12.1 | 52.7 | R | Cerebelum (IX) | 1.7 | 4.38 | 8, -44, -42 |
| 21 |  |  | R | Superior Frontal Gyrus |  | 4.05 | 28, 2, 68 |

This Table represents a summary of the main results. Brain regions are referred to as by the nomenclature of the Anatomy Toolbox (Eickhoff et al., 2005). Only main peaks are reported. Clusters with voxels in cyto < 20 were not reported. Clusters with N/A areas only where not reported. Full results are displayed in a .txt file entitled TABLE S2.A on OSF.

**TABLE S2.B - VICTIM FIRST - Regression  [FreeShocks-NoShock] - vindictiveness,  *negative relationship, FWE corrected at cluster size***

| ***cluster size (N voxels)*** | **#Voxels in cyto** | **% Cluster** | **Hem** | **Cyto or Anatomical description** | **%**  **Area** | **cluster peak t-value** | **MNI coordinates (X,Y,Z)** |
| --- | --- | --- | --- | --- | --- | --- | --- |
| 949 | 113.8 | 40.6 | L | Lingual Gyrus | 12 | 6.48 | -6, -68, 4 |
|  | 83.3 | 8.8 | L | Cerebelum (VI) | 4.4 | 5.91 | -4, -68, -12 |
|  | 385.5 | 40.6 | L | Lingual Gyrus | 19.1 | 5.38 | 0, -74, 2 |
|  | 385.5 | 40.6 | L | Lingual Gyrus | 19.1 | 5.36 | -16, -78, 2 |
|  | 14.6 | 1.5 |  | Cerebellar Vermis (4 / 5) | 1.8 | 5.10 | 0, -58, 0 |
|  | 385.5 | 40.6 | L | Lingual Gyrus | 19.1 | 5.07 | -12, -58, -2 |
|  | 385.5 | 40.6 | L | Lingual Gyrus | 19.1 | 5.06 | -10, -58, -2 |
|  | 42 | 4.4 |  | Cerebellar Vermis (4 / 5) | 2 | 4.92 | 4, -60, 2 |
|  |  |  | L | Calcarine Gyrus |  | 4.83 | -16, -62, ,8 |
|  | 385.5 | 40.6 | L | Calcarine Gyrus | 19.1 | 4.29 | -16, -78, 12 |
|  | 113.8 | 40.6 | L | Calcarine Gyrus | 12 | 4.28 | -22, -68, 8 |
| 698 |  |  | L | Precuneus |  | 7.58 | -6, -62, 48 |
|  |  |  |  | Precuneus |  | 7.53 | -10, -58, 50 |
|  |  |  | L | Precuneus |  | 6.91 | -4, -58, 48 |
|  |  |  |  | N/A |  | 6.87 | -16, -54, 46 |
|  |  |  | L | Precuneus |  | 6.36 | -4, -58, 54 |
|  | 61.6 | 8.8 | L | Precuneus | 17.7 | 6.04 | -6, -72, 56 |
|  | 61.6 | 8.8 | L | Precuneus | 17.7 | 5.84 | -10, -74, 56 |
|  | 99.9 | 14.3 | L | Superior Parietal Lobule | 8 | 5.68 | -14, -74, 54 |
|  | 61.6 | 8.8 | L | Superior Occipital Gyrus | 17.7 | 5.11 | -6, -80, 42 |
|  | 1.3 | 0.2 | L | Precuneus | 0.2 | 4.37 | -4, -50, 68 |
|  |  |  | L | Superior Parietal Lobule |  | 4.14 | -14, -70, 48 |
| 502 | 21.9 | 4.4 | L | Hippocampus | 10 | 7.27 | -36, -28, -12 |
|  | 74.1 | 14.8 | L | Thalamus | 23.3 | 6.52 | -18, -26, 4 |
|  | 74.1 | 14.8 | L | Thalamus | 23.3 | 6.11 | -22, -28, 6 |
|  |  |  |  | N/A |  | 5.82 | -24, -30, 8 |
|  |  |  | L | ParaHippocampal Gyrus |  | 5.52 | -28, -36, -12 |
|  | 6.3 | 1.2 | L | Hippocampus | 15.5 | 5.35 | -30, -20, -14 |
|  |  |  | L | N/A |  | 5.29 | -32, -32, 10 |
|  |  |  | L | Hippocampus |  | 5.27 | -36, -18, -14 |
|  |  |  | L | Fusiform Gyrus |  | 5.23 | -38, -16, -20 |
|  | 29.9 | 6 | L | ParaHippocampal Gyrus | 18.2 | 5.10 | -26, -38, -6 |
|  | 16.4 | 3.3 | L | Fusiform Gyrus | 2 | 5.01 | -42, -44, -20 |

Only clusters surviving a 5% FWE correction at the cluster size are reported (t=3.79, *p* < .001, cluster size 165). Brain regions are referred to as by the nomenclature of the Anatomy Toolbox (Eickhoff et al., 2005). The file is downloadable as a .txt file entitled ‘Table S2.B’ on OSF.

SUPPLEMENTARY FIGURE S3

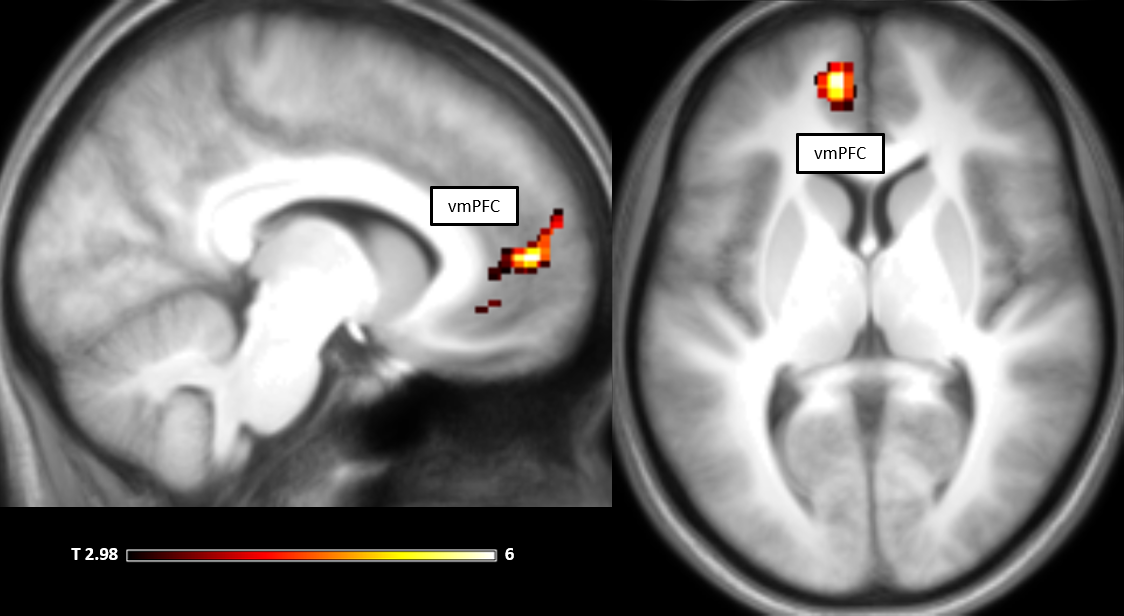

**Fig. S3.** **MRI results** [FreeDecisionPhase - CoercedDecisionPhase] contrast. Peak coordinates can be seen in Table S3.  Results are corrected using 5% FWE at cluster level (280 voxels), t=3.37, *p* < .001 at voxel level). Peak coordinates of the cluster surviving FWE correction at the cluster size can be seen in Table S3. The reverse relationship was not significant (for a t=3.37, *p* < .001).

SUPPLEMENTARY TABLE S3

**TABLE S3 - [FreeDecisionPhase - CoercedDecisionPhase] contrast**

| ***cluster size (N voxels)*** | **#Voxels in cyto** | **% Cluster** | **Hem** | **Cyto or Anatomical description** | **%**  **Area** | **cluster peak t-value** | **MNI coordinates (X,Y,Z)** |
| --- | --- | --- | --- | --- | --- | --- | --- |
| 280 | 40.3 | 14.4 | L | Area Fp2 - Superior Medial Gyrus | 5.6 | 4.61 | -6, 58, 14 |
|  | 15 | 5.4 | L | Area s32 - Mid Orbital Gyrus | 7.2 | 3.97 | -10, 40, -8 |
|  |  |  | L | Area s32 - ACC |  | 3.93 | -14, 46, 0 |
|  |  |  | L | Area s32 - ACC |  | 3.92 | -6, 42, 0 |
|  | 3.6 | 1.3 | L | Area Fp1 - Superior Frontal Gyrus | 0.2 | 4.57 | -20, 54, 0 |
|  |  |  | L | Area Fp1 - Mid Orbital Gyrus |  | 4.04 | -12, 42, -6 |
|  | 1.1 | 0.4 | L | Area s24 - Mid Orbital Gyrus | 0.7 | 3.45 | -4, 34, -12 |

Only clusters surviving a 5% FWE correction at the cluster size are reported (t=3.37, *p* < .001, cluster size 280). Brain regions are referred to as by the nomenclature of the Anatomy Toolbox (Eickhoff et al., 2005).
